# Learning unsupervised feature representations for single cell microscopy images with paired cell inpainting

**DOI:** 10.1101/395954

**Authors:** Alex X Lu, Oren Z Kraus, Sam Cooper, Alan M Moses

## Abstract

Cellular microscopy images contain rich insights about biology. To extract this information, researchers use features, or measurements of the patterns of interest in the images. Here, we introduce a convolutional neural network (CNN) to automatically design features for fluorescence microscopy. We use a self-supervised method to learn feature representations of single cells in microscopy images without labelled training data. We train CNNs on a simple task that leverages the inherent structure of microscopy images and controls for variation in cell morphology and imaging: given one cell from an image, the CNN is asked to predict the fluorescence pattern in a second different cell from the same image. We show that our method learns high-quality features that describe protein expression patterns in single cells both yeast and human microscopy datasets. Moreover, we demonstrate that our features are useful for exploratory biological analysis, by capturing high-resolution cellular components in a proteome-wide cluster analysis of human proteins, and by quantifying multi-localized proteins and single-cell variability. We believe paired cell inpainting is a generalizable method to obtain feature representations of single cells in multichannel microscopy images.

**Author Summary:** To understand the cell biology captured by microscopy images, researchers use features, or measurements of relevant properties of cells, such as the shape or size of cells, or the intensity of fluorescent markers. Features are the starting point of most image analysis pipelines, so their quality in representing cells is fundamental to the success of an analysis. Classically, researchers have relied on features manually defined by imaging experts. In contrast, deep learning techniques based on convolutional neural networks (CNNs) automatically learn features, which can outperform manually-defined features at image analysis tasks. However, most CNN methods require large manually-annotated training datasets to learn useful features, limiting their practical application. Here, we developed a new CNN method that learns high-quality features for single cells in microscopy images, without the need for any labeled training data. We show that our features surpass other comparable features in identifying protein localization from images, and that our method can generalize to diverse datasets. By exploiting our method, researchers will be able to automatically obtain high-quality features customized to their own image datasets, facilitating many downstream analyses, as we highlight by demonstrating many possible use cases of our features in this study.

## Introduction

Feature representations of cells within microscopy images are critical for quantifying cell biology in an objective way. Classically, researchers have manually designed features that measure phenomena of interest within images: for example, a researcher studying protein subcellular localization may measure the distance of a fluorescently-tagged protein from the edge of the cell (1), or the correlation of punctuate proteins with microtubules (2). By extracting a range of different features, an image of a cell can be represented as a set of values: these feature representations can then be used for numerous downstream applications, such as classifying the effects of pharmaceuticals on cancer cells (3), or exploratory analyses of protein localization (1,4). The success of these applications depends highly on the quality of the features used: good features are challenging to define, as they must be sensitive to differences in biology, but robust to nuisance variation such as microscopy illumination effects or single cell variation (5).

Convolutional neural networks (CNNs) have achieved state-of-the-art performance in tasks such as classifying cell biology in high-content microscopy and imaging flow cytometry screens (6–9), or segmenting single cells in images (10,11). A key property driving this performance is that CNNs automatically learn features that are optimized to represent the components of an image necessary for solving a training task (12). Donahue *et al.* (13) and Razavian *et al.* (14) demonstrate that the feature representations extracted by the internal layers of CNNs trained as classifiers achieve state-of-the-art results on even very different applications than the task the CNN was originally trained for; studies specific to bio-imaging report similar observations about the quality of CNN features (6,15,16). The features learned by CNNs are thought to be more sensitive to relevant image content than human-designed features, offering a promising alternative for feature-based image analysis applications.

However, learning high-quality features with CNNs is a current research challenge because it depends highly on the training task. For example, autoencoders, or unsupervised CNNs trained to reconstruct input images, usually do not learn features that generalize well to other tasks (17–19). On the other hand, classification tasks result in high-quality features (13,14), but they rely on large, manually-labeled datasets, which are expensive and time-consuming to generate. For example, to address the challenge of labeling microscopy images in the Human Protein Atlas, the project launched a massive crowd-sourcing initiative in collaboration with an online video game, spanning over one year and involving 322,006 gamers (20). Because advances in high-content throughput microscopy are leading to routine generation of thousands of images (21), the new cell morphologies and phenotypes discovered and need for integration of datasets (4) may require the continuous update of models. Unsupervised methods that result in the learning of high-quality features without the use of manual labels would resolve the bottleneck of collecting and updating labels: we would, in principle, be able to learn feature representations for any dataset, without the need for experts to curate, label, and maintain training images.

If obtaining expert-assigned labels for microscopy images is challenging, then obtaining labels for the single cells within these images is even more difficult. Studying biological phenomena at a single-cell level is of current research interest (22): even in genetically-identical cell cultures, single-cell variability can originate from a number of important regulatory mechanisms (23), including the cell cycle (24), stochastic shuttling of transcription factors between compartments (25), or variability in the DNA damage response (26). Thus, ideally, a method would efficiently generate feature representations of single cells for arbitrary datasets using deep learning, without the need for labelled single-cell training data.

In this study, we asked if CNNs could automatically learn high-quality features for representing single cells in microscopy images, without using manual labels for training. We investigate *self-supervised learning,* which proposes training CNNs using labels that are automatically available from input images. Self-supervised learning aims to develop a feature representation in the CNN that is useful for other tasks: the training task is only a pretext, and the CNN may not achieve a useful level of performance at this pretext task. This differs from weakly-supervised learning, where a learnt network is used directly for an auxiliary task (27), such as segmenting tumors in histology images using a network trained to predict disease class (28). After training, the output of the CNN is discarded, and internal layers of the CNN are used as the features. The logic is that by learning to solve the pretext task, the CNN will develop features that are useful for other applications.

The central challenge in self-supervised learning is defining a pretext task that encourages the learning of generalizable features (29). Successful self-supervised learning strategies in the context of natural images, include CNNs trained to predict the appearance of withheld image patches based upon its context (18), the presence and location of synthetic artifacts in images (29), or geometric rotations applied to input images (30). The idea is that to succeed at the pretext task, the CNN needs to develop a strong internal representation of the objects and patterns in the images. When transferred to tasks such as classification, segmentation, or detection, features developed by self-supervised methods have shown state-of-the-art results compared to other unsupervised methods, and in some cases, perform competitively with features learned by supervised CNNs (17,29–32).

Here, we present a novel self-supervised learning method designed for microscopy images of protein expression in single cells. Our approach leverages the typical structure of these images to define the pretext training task: in many cases, each image contains multiple genetically identical cells, growing under the same experimental condition, and these cells exhibit similar patterns of protein expression. The cells are imaged in multiple “channels” (such as multiple fluorescence colours) that contain very different information. By exploiting this structure, we define a pretext task that relies only upon image content, with no prior human labels or annotations incorporated. In our examples, one set of channels represents one or more structures in the cell (e.g. the cytosol, nucleus, or cytoskeleton), and another channel represents proteins of interest that have been fluorescently tagged to visualize their localization (with a different protein tagged in each image). Then, given both channels for one cell and the structural markers for a different cell from the same image, our CNN is trained to predict the appearance of the protein of interest in the second cell, a pretext task that we term “paired cell inpainting” (Figure 1A). To solve this pretext task, we reason that the CNN must identify protein localization in the first (or “source”) cell and reconstruct a similar protein localization in the second (or “target”) cell in a way that adapts to single cell variability. In Figure 1, the protein is localized to the nucleoli of human cells – the network must recognize the localization of the protein in the source cell, but also transfer it to the equivalent structures in the target cell, despite differences in the morphology of the nucleus between the two cells. Thus, by the design of our pretext task, our method learns representations of single cells.

**Fig 1.**
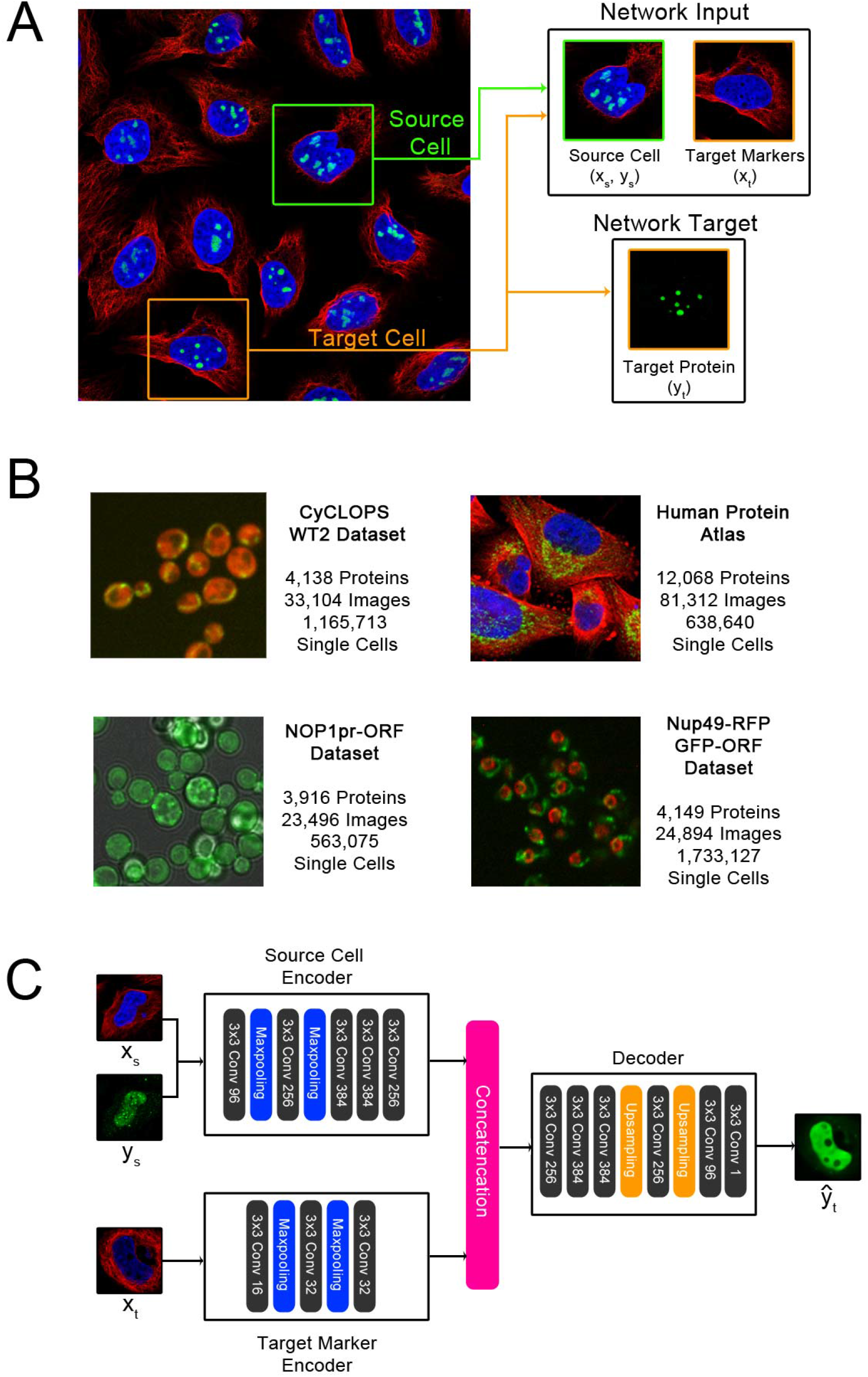
An overview of the inputs and targets to the network in paired cell inpainting, and of our proposed architecture. (A) Inputs and targets to the network. We crop a source cell (green border) and a target cell (orange border) from the same image. Then, given all channels for the source cell, and the structural markers for the target cell (in this dataset, the nucleus and the microtubule channels), the network is trained to predict the appearance of the protein channel in the target cell. Images shown are of human cells, with the nucleus colored blue, microtubules colored red, and a specific protein colored green. (B) Example images from the proteome-scale datasets we use in this study. We color the protein channel for each dataset in green. The CyCLOPS yeast dataset has a cytosolic RFP, colored in red. The NOP1pr-ORF yeast dataset has a brightfield channel, colored grey. The Nup49-RFP GFP-ORF yeast dataset has a RFP fused to a nuclear pore marker, colored in red. The Human Protein Atlas images are shown as described above in A. (C) Our proposed architecture. Our architecture consists of a source cell encoder and a target marker encoder. The final layers of both encoders are concatenated and fed into a decoder that outputs the prediction of the target protein 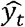. Layers in our CNN are shown as solid-colored boxes; we label each of the convolutional layers (solid grey boxes) with the number of filters used (e.g. 3×3 Conv 96 means 96 filters are used). We show a real example of a prediction from our trained human cell model given the input image patches in this schematic.

To demonstrate the generalizability of our method, we automatically learn feature representations for datasets of both human and yeast cells, with different morphologies, microscopes and resolutions, and fluorescent tagging schemes (Figure 1B). We exemplify the quality of the features learned by our models in several use-cases. First, we show that for building classifiers, the features learned through paired cell inpainting improve in discriminating protein subcellular localization classes at a single-cell level compared to other unsupervised methods. Next, we establish that our features can be used for unsupervised exploratory analysis, by performing an unsupervised proteome-wide cluster analysis of protein analysis in human cells, capturing clusters of proteins in cellular components at a resolution challenging to annotate by human eye. Finally, we determine that our features are useful for single-cell analyses, showing that our features can distinguish phenotypes in spatially-variable single cell populations.

## Methods

### Paired Cell Inpainting

We would like to learn a representation for single cells in a collection of microscopy images, *I*. We define each image *i* as a collection of single cells, *i* = {*c*_*i*,1_ …*c*_*i,n*_}. We note that the only constraint on *i* is that its single cells *C*_*i*_ must be considered similar to each other, so *i* does not need to be strictly defined as a single digital image so long as this is satisfied; in our experiments, we consider an “image” *i* to be all fields of view corresponding to an experimental well.

We define single cells to be image patches, so *c* ∈ ℝ ^*H*×*W*×*Z*^, where *Z* are the channels. We split the images by channel into *c* = (*x, y*), where 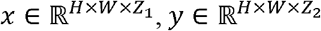, and *Z*_1_, *Z*_2_ ⊆ *Z*. For this work, we assign to *Z*_1_ channels corresponding to structural markers, or fluorescent tags designed to visualize structural components of the cell, where all cells in the collection of images have been labeled with the tag. We assign to *Z*_2_ channels corresponding to proteins, or channels where the tagged biomolecule will vary from image to image.

We define a source cell *c*_*s*_, which is associated with a target cell *c*_*t*_ satisfying constraints that both cells are from the same image, *c*_*s*_ ∈ *i*_*s*_, *c*_*t*_ ∈ *i*_*t*_, *i*_*s*_ = *i*_*t*_, and *c*_*s*_ ≠ *c*_*t*_. Our goal is to train a neural network that will solve the prediction problem 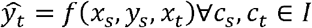, where 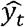 represents the predicted protein channels that vary between images.

For this work, we train the network on the prediction problem by minimizing a standard pixel-wise mean-squared error loss between the predicted target protein 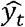 and the actual target protein *y*_*t*_:

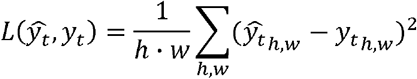

As with other self-supervised methods, our pretext training task is only meant to develop the internal feature representation of the CNN. After training, 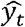 is discarded, and the CNN is used as a feature extractor. Importantly, while our pretext task predicts a label *y*_*t*_, we consider our overall method to be unsupervised, because these labels are defined automatically from image content without any human supervision.

One limitation in our pretext task is that some protein localizations are not fully deterministic in respect to the structure of the cell and are therefore challenging to predict given the inputs we define. In these cases, we observe that the network produces smoothed predictions that we hypothesize are an averaged guess of the localization. We show an example in Figure 1C; while the source protein is localized to the nucleus in a punctate pattern, the target protein is predicted as a smooth distribution throughout the nucleoplasm. However, as our inpainting task is a pretext and discarded after training, we are not concerned with outputting fully realistic images. The averaging effect is likely due to our choice of a mean squared loss function; should more realistic images be desired, different loss functions, such as adversarial losses (33), may produce better results.

### Architecture

As the goal of our training task is to obtain a CNN that can encode single cell image patches into feature representations, we construct independent encoders for(*x*_*s*_, *y*_*s*_) and for *x*_*t*_, which we call the “source cell encoder” and the “target marker encoder”, respectively. After training with our self-supervised task, we isolate the source cell encoder, and discard all other components of our model. This architecture allows us to obtain a feature representation of any single cell image patch independently without also having to input target cell markers. To obtain single cell features, we simply input a single cell image patch, and extract the output of an intermediate convolutional layer in the source cell encoder.

We show a summary of our architecture in Figure 1C. Following other work in self-supervised learning (17,18,30), we use an AlexNet architecture for the source cell encoder, although we set all kernel sizes to 3 due to the smaller sizes of our image patches, and we add batch normalization after each convolutional layer. We use a smaller number of filters and fewer convolutional layers in the architecture of the target marker encoder; we use three convolutional layers, with 16, 32, and 32 filters, respectively. Finally, for the decoder, we reverse the AlexNet architecture.

The goal of our training is to develop the learned features of our source cell encoder, which will later be used to extract single cell feature representations. If the network were to utilize bleed-through from the fluorescent marker channels to predict the target marker image, or overfit to the data and ‘memorize’ the target cell images, then the network would not need to learn to extract useful information from the source cell images. To rule out that our trained models were subject to these effects, we produced inpainting results from a trained model, where we paired source cells with target cells where there is a mismatch between source and target protein localization. Here, the model has seen both the source and target cells during training, but never the specific pair due to the structure of our training task, as the cells originate from different images. We qualitatively confirmed that our trained networks were capable of synthesizing realistic results agreeing with the protein localization of the source cell (Supplementary Figure 1), suggesting that our models are not trivially overfitting to the target marker images.

### Sampling Pairs

Because the training inputs to our self-supervised task consist of pairs of cells, the number of possible combinations is large, as each cell may be paired with one of many other cells. This property is analogous to data augmentation, increasing the number of unique training inputs to our network. To sample training inputs, for each epoch, we iterate through every single cell in our training dataset, set it as the source cell, and draw with uniform probability a target cell from the set of all valid possible target cells.

Our pretext task relies on the assumption that protein expression in single cells from the same image is similar. This assumption is not always true: in the datasets used in this work, some proteins exhibit significant single cell variability in their protein abundance or localization (34,35). These proteins may contribute noise because paired single cells will have (unpredictably) different protein expression patterns and the model will not learn. Although the Human Protein Atlas documents variable proteins (24), for our experiments we do not remove these and confirm that our model still learns good features in spite of this noise.

### Datasets, Preprocessing, and Training

For yeast cells, we used the WT2 dataset from the CYCLoPS database (36). This collection expresses a cytosolic red fluorescent protein (RFP) in all cells, and tags proteins of interest with green fluorescent protein (GFP). We use the RFP channel as the structural marker, and the GFP channel as the protein. To extract single cell crops from the images in this dataset, we segmented our images using YeastSpotter on the structural marker channel (37), and extract a 64×64 pixel crop around the identified cell centers; we discarded any single cells with an area smaller than 5% or greater than 95% of the image crop, as these are likely artifacts arising from under- or over-segmentation. We discard any images with fewer than 30 cells. We preprocessed crops by rescaling each crop to be in the range of [0, 1]. These preprocessing operations result in a total of 1,165,713 single cell image patches grouped into 4,069 images (where each image is 4 fields of view), with a total of 4,069 of 4,138 proteins passing this filter.

We also trained a second different yeast cell model, using a NOP1pr-GFP library previously published by Weill *et al.* (37) This collection tags all proteins of interest with GFP at the N-terminal of proteins with a NOP1 promoter, and is also imaged in brightfield. We used the brightfield channel as a structural marker and the GFP channel as the protein. We extracted and preprocessed single cell crops using the same procedure for the CYCLoPS dataset, and discarded any images with fewer than 30 single cells. These preprocessing operations result in a total of 563,075 single cell image patches grouped into 3,067 images (where each image is 3 fields of view), with a total of 3,067 of 3,916 proteins passing this filter.

Finally, we trained a third yeast cell model, using a dataset previously published by Tkach *et al.* (38), a Nup49-RFP GFP-ORF library. This collection expresses a nuclear pore protein (Nup49) fused to RFP in all cells, and tags proteins of interest with green fluorescent protein (GFP). We use the RFP channel as the structural marker, and the GFP channel as the protein. We extracted and preprocessed single cell crops using the same procedure for the CYCLoPS dataset, and discarded any images with fewer than 30 single cells. These preprocessing operations result in a total of 1,733,127 single cell image patches grouped into 4,085 images (where each image is 3 fields of view), with a total of 4,085 of 4,149 proteins passing this filter.

For human cells, we use images from version 18 of the Human Protein Atlas (24). We were able to download jpeg images for a total of 12,068 protein. Each protein may have multiple experiments, which image different cell line and antibody combinations. We consider an image to be of a protein for the same cell line and antibody combination; accordingly, we have 41,517 images (where each image is usually 2 fields of view). We downloaded 3 channels for these images. Two visualize the nuclei and microtubules, which we use as the structural marker channels. The third channel is an antibody for the protein of interest, which we use as the protein channel. To extract single cell crops from this image, we binarize the nuclear channel with an Otsu filter and find nuclei by labeling connected components as objects using the scikit-image package (39). We filter any objects with an area of less than 400 pixels, and extract a 512×512 pixel crop around the center of mass of remaining objects. To reduce training time, we rescale the size of each crop to 64×64 pixels. We preprocessed crops by rescaling each crop to be in the range of [0, 1], and clipped pixels under 0.05 intensity for the microtubule and nuclei channels to 0 to improve contrast. Finally, we remove any images with fewer than 5 cells, leaving a total of 638,640 single cell image patches grouped into 41,285 images, with a total of 11,995 of 12,068 proteins passing this filter.

Crop sizes for our datasets (64×64 pixels for yeast cells and 512×512 pixels for human cells) were chosen such that a crop fully encompasses an average cell from each of these datasets. We note that different image datasets may require different crop sizes, depending on the resolution of the images and the size of the cells. While each crop is centered around a segmentation, we did not filter crops with overlapping or clumped cells, so some crops may contain multiple cells. In general, we observed that our models did not have an issue learning to inpaint protein expression from a crop with multiple cells to a crop with a single cell, or vice versa: Supplementary Figure 1 shows an example of a case where we inpaint protein expression from a crop with two cells to a crop with one cell.

During training, we apply random horizontal and vertical flips to source and target cells independently as data augmentation. We trained models for 30 epochs using an Adam optimizer with an initial learning rate of 1e-4.

After training, we extract representations by maximum pooling the output of an immediate convolutional layer, across spatial dimensions. This strategy follows previous unsupervised representation extraction from self-supervised learning methods (17,30), which sample activations from feature maps.

### Baseline Feature Extraction and Benchmarking

To benchmark the performance of features learned using paired cell painting, we obtained features from other commonly-used feature representation strategies.

As classic computer vision baselines for our yeast cell benchmarks, we obtained features extracted using CellProfiler (40) for a classification dataset of 30,889 image crops of yeast cells directly from the authors (41). These features include measurements of intensity, shape, and texture, and have been optimized for classification performance. Further details are available from (41). We also extracted features from these yeast cell image crops using interpretable expert-designed features by Handfield *et al.* (1) We followed procedures previously established by the authors: we segmented cells using provided software, and calculated features from the center cell in each crop.

For our transfer learning baselines in our yeast cell benchmarks, we used a VGG16 model pretrained on ImageNet, using the Keras package. We benchmarked three different input strategies: (1) we mapped channels arbitrarily to RGB channels (RFP to red, GFP to green, blue channel left empty); (2) we inputted each channel separately as a greyscale image and concatenated the representations; (3) we inputted only the GFP channel as a greyscale image and used this representation alone. In addition, we benchmarked representations from each convolutional layer of VGG16, post-processed by maximum pooling across spatial dimensions (as we did for our self-supervised features). Supplementary Figure 2 shows classification performance using each layer in VGG16 with the *k*NN classifier described in our benchmarks, across all three strategies. In general, we observed that the 3^rd^ input strategy resulted in superior performance, with performance peaking in the 4^th^ convolutional block of VGG16. We report results from the top-performing layer using the top-performing input strategy in our benchmarks.

Contrary to previous work in transfer learning on microscopy images by Pawlowski *et al.*, we extract features from the intermediate convolutional layers of our transferred VGG16 model instead of the final fully-connected layer (15). This modification allows us to input our images at their original size, instead of resizing them to the size of the images originally used to train the transferred model. As our work operates on single cell crops, which are much smaller than the full images benchmarked in previous work (64×64 pixels compared to 1280×1024 pixels), we found that inputting images at their original size instead of stretching them resulted in performance gains: our top-performing convolutional layer (block4_conv1) with inputs at original size achieved 69.33% accuracy, whereas resizing the images to 224×224 and using features from the final fully-connected layer (as-is, without max-pooling, as described in (15)) achieves 65.64% accuracy. In addition, we found that extracting features from images at original resolution improves run-time: while inputting resized 224×224 images and extracting features from the final fully-connected layer took 0.31 seconds per image on average, inputting 64×64 crops and extracting features from the best-performing layer only took 0.02 seconds per image.

Finally, for the supervised baseline, we used the model and pretrained weights provided by Kraus *et al.* (6). We inputted images as previously described. To ensure that metrics reported for these features were comparable with the other accuracies we reported, we extracted features from this model and built the same classifier used for the other feature sets. We found that features from the final fully connected layer before the classification layer performed the best, and report results from this layer.

To compare the performance of various feature representations with our single yeast cell dataset, we built *k*NN classifiers. We preprocessed each dataset by centering and scaling features by their mean and standard deviation. We employed leave-one-out cross-validation and predicted the label of each cell based on its neighbors using Euclidean distance. Supplementary Table 1 shows classification accuracy with various parameterizations of *k*. We observed that regardless of *k*, feature representations were ranked the same in their classification performance, with our paired cell inpainting features always outperforming other unsupervised feature sets. However, *k* = 11 produced the best results for all feature sets, so we report results for this parameterization.

As classic computer vision baselines for our human cell benchmarks, we curated a set of texture, correlation, and intensity features. For each crop, we measured the sum, mean, and standard deviation of intensity from pixels in the protein channels, and the Pearson correlation between the protein channel and the microtubule and nucleus channels. We extracted Haralick texture features from the protein channel at 5 scales (1, 2, 4, 8, and 16 pixels).

Finally, as the transfer learning baseline for our human cell benchmarks, we extracted features from the pretrained VGG16 model using the same input strategies and layer as established in our yeast cell benchmark.

### Comparing Pairwise Distances in Feature Spaces

To directly measure and compare how a feature set groups together cells with similar localizations in their feature spaces, measured the average pairwise distance between cells in the feature space. We preprocess single cell features by scaling to zero mean and unit variance, to control for feature-to-feature differences in scaling within feature sets. Then, to calculate the distance between two single cells *c*, we use the Euclidean distance between their features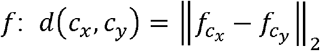.

Given two images with the same localization term, we calculate the average distance of all cells in the first image paired with all cells in the second image and normalize these distances to an expectation of the average pairwise distance between images with different localization terms. A negative normalized average pairwise distance score indicates that the distances are smaller than expectation (so single cells in images with the same label are on average closer in the feature space).

For the Human Protein Atlas images, we restricted our analysis to proteins that only had a single localization term shared by at least 30 proteins, resulting in proteins with 18 distinct localization terms (as listed in Figure 2B). For each localization term, we calculated average pairwise distances for 1,000 random protein pairs with the same localization term, relative to an expectation from 1,000 random protein pairs with each of the possible other localization terms (for a total of 17,000 pairs sampled to control for class imbalance). For our experiments controlling for cell line, we also introduce the constraint that the images must be of cells of the same or different cell lines, depending on the experiment. Because some localization terms and cell line combinations are rare, we did not control for class imbalance and drew 10,000 random protein pairs with any different localization terms (not necessarily each other different term), and compared this to 10,000 random protein pairs with the same localization term. Hence, the distances in the two experiments are not directly comparable.

**Fig 2.**
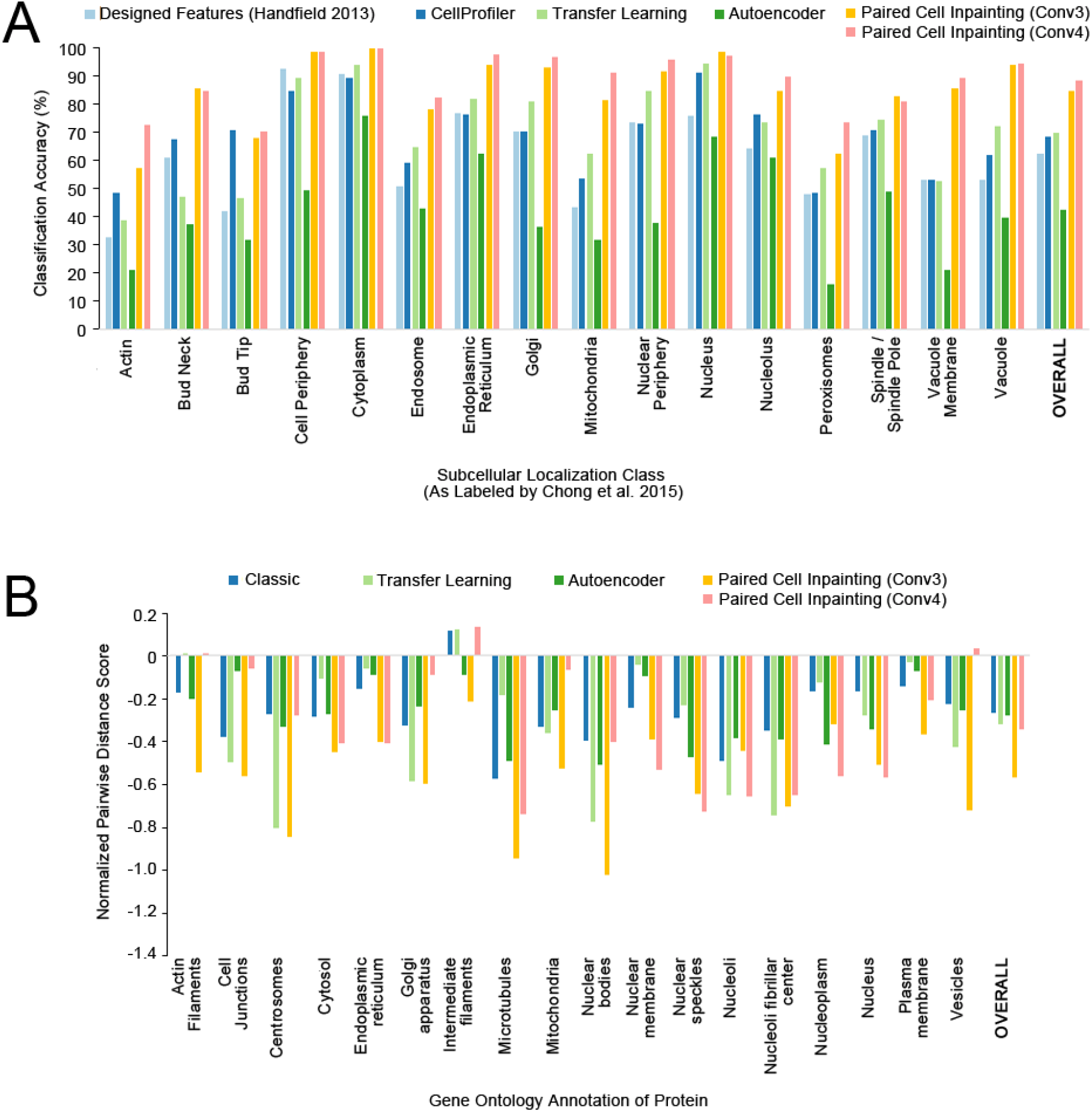
Quantitative comparisons of paired cell inpainting features with other unsupervised feature sets. (A) Overall and class-by-class performance benchmark for yeast single cell protein localization classes using unsupervised feature sets and a *k*NN classifier (*k* = 11) on our test set. For all approaches extracting features from CNNs (Transfer Learning, Autoencoder, and Paired Cell Inpainting), we extract representations by maximum pooling across spatial dimensions, and report the top-performing layer. We report the overall accuracy as the balanced accuracy of all classes. (B) The normalized pairwise distance between cells with the same localization terms according to gene ontology labels, for proteins in the Human Protein Atlas. A more negative score indicates that cells are closer in the feature set compared to a random expectation.

### Single cell analysis of Multiply-Localized Proteins

For proteins localized to two compartments, we calculated a score for each cell based upon its distance to the first compartment versus its distance to the second compartment. To do so, we averaged the feature vectors for all single-cells in images annotated to localize to each compartment alone to define the average features for the two compartments. Then, for every single-cell, we calculated the distance of the single-cell’s features relative to the two compartments’ averages and take a log ratio of the distance to the first compartment divided by the distance to the second compartment. A negative number reflects that the single-cell is closer to the first compartment in the feature space, while a positive number reflects that the single-cell is closer to the second compartment.

## Experimental Results

### Paired cell inpainting features discriminate protein subcellular localization in yeast single cells

To assess the quality of feature learned through paired cell inpainting we trained a model for yeast fluorescent microscopy images using paired cell inpainting, on an unlabelled training set comprising of the entire WT2 screen in the CyCLOPS dataset, encompassing 1,165,713 single cells from 4,069 images.

Good features are sensitive to differences in biology, but robust to nuisance variation (5). As a first step to understanding if our features had these properties, we compared paired cell inpainting features with other feature sets at the task of discriminating different subcellular localization classes in yeast single cells. To do this, we made use of a test set of 30,889 single cell image patches manually assigned to 17 different protein subcellular localization classes by Chong *et al.* (41). These single cells have been curated from a different image screen than the one we used to train our model, and thus represent an independent test dataset that was never seen by the model during training.

To evaluate feature sets, we constructed a simple *k*NN classifier (*k* = 11) and evaluated classification performance by comparing the predicted label of each single cell based on its neighbors to its actual label. While more elaborate classifiers could be used, the *k*NN classifier is simple and transparent, and is frequently employed to compare feature sets for morphological profiling (15,16,42,43). Like these previous works, the goal of our experiment is to assess the relative performance of various feature sets in a controlled setting, not to present an optimal classifier.

To use the feature representation from self-supervised CNN, we must first identify which layers represent generalizable information about protein expression patterns. Different layers in self-supervised models may have different properties: the earlier layers may only extract low-level features, but the later layers may be too specific to production of the pretext task (12). For this reason, identifying self-supervised features with a layer-by-layer evaluation of the model’s properties is standard (18–20,28,29,31). Since our model has five convolutional layers, it can be interpreted as outputting five different feature sets for each input image. To determine which feature set would be best for the task of classifying yeast single cells, we constructed a classifier on our test set for the features from each layer independently, as shown in Table 1. Overall, we observe that the third (Conv3) and fourth (Conv4) convolutional layers result in the best performance.

**Table 1.** Classification accuracies for various layers in our CNN.

| **Layer** | **Overall Classification Accuracy** |
| --- | --- |
| **Conv1** | 43.09 |
| **Conv2** | 70.52 |
| **Conv3** | **84.30** |
| **Conv4** | **87.98** |
| **Conv5** | 81.89 |

Classification accuracies using various layers in our CNN, for single yeast cell localization classes using a *k*NN classifier (*k* = 11) on our test set. For each layer, we extract a representation by maximum pooling feature maps across spatial dimensions. We report the overall accuracy as the balanced accuracy of all classes.

Next, we sought to compare our features from paired cell painting with other unsupervised feature sets commonly used for cellular microscopy images. First, we compared our features with feature sets designed by experts: CellProfiler features (40) that were previously extracted from segmented single cells by Chong *et al.* to train their single cell classifier (41), and with interpretable features measuring yeast protein localization calculated using yeast-specific segmentation and feature extraction software developed by Handfield *et al.* (1). Second, we compared our features with those obtained from transfer learning, i.e., repurposing features from previously-trained supervised CNNs in other domains (13,14): we used features from a VGG16 model (44), a previous supervised CNN trained on Imagenet, a large-scale classification dataset of natural images (45). Finally, we transferred features from an autoencoder (an unsupervised CNN) trained on the same dataset as our paired cell inpainting models, using our source cell encoder and decoder architecture.

Overall, the CellProfiler and VGG16 features perform at 68.15% and 69.33% accuracy respectively on our evaluation set, while the yeast-specific, interpretable features are worse at 62.04%. Autoencoder features achieve 42.50% accuracy. Our features from convolutional layer 4 (Conv4) perform better than any of the other features benchmarked by a large margin, achieving 87.98% (Figure 2A).

As an estimate of the upper bound of classification performance with our *k*NN classifier, we evaluated the performance of features extracted by the layers of the state-of-the-art supervised CNN single-cell classifier trained with most of the test set used here (6). The features from the best performing layer from this supervised model achieve 92.39% accuracy, which is comparable to the 87.98% accuracy achieved by the self-supervised paired cell inpainting features. Taken together, these results suggest that paired cell inpainting learns features comparable with those learned by supervised neural networks for discriminating protein subcellular localization in yeast single cells.

To test if these improvements in classification performance were agnostic to the kind of classifier used, we also benchmarked the performance of logistic regression and random forest classifiers built with all feature sets (Supplementary Table 2). We observed that our paired cell inpainting features continue to outperform other unsupervised feature sets by a large margin, even with different classification models.

To assess if the improvements in classification performance over other unsupervised methods was because our features form a general global representation of protein localization, as opposed to just clustering similar cells locally, we visualized features for this yeast single cell dataset using UMAP (46) (Supplementary Figure 3). We observed that with our paired cell inpainting features, single cells with the same label cluster together in a continuous distribution in the UMAP-reduced space, similar to those learned by the fully-supervised model, whereas this effect is not as strong with CellProfiler features or VGG16 features.

Finally, to assess if our method could learn effective representations for yeast image datasets regardless of modality or fluorescent tagging scheme, we trained two new models on two additional datasets: a yeast GFP-ORF collection imaged in brightfield (47), and a yeast GFP-ORF collection imaged with a nuclear pore structural marker (38). We averaged the features of all single cells for each protein for each model and visualized the protein representations using UMAP (shown in Figure 3, along with example images from each dataset), coloring points using previous manual labels on the protein level (47,48). We observed distinct clusters for most protein localization labels for the representation produced by each model. Remarkably, even though each model is independent and trained on an independent dataset, we observed a substantial degree of consistency between clusters in the distances of their learned representations: for example, in all three representations, the “nucleolus” cluster was close to the “nucleus” cluster, and the “ER” and “cytoplasm and nucleus” clusters were adjacent to the “cytoplasm” cluster.

**Fig 3.**
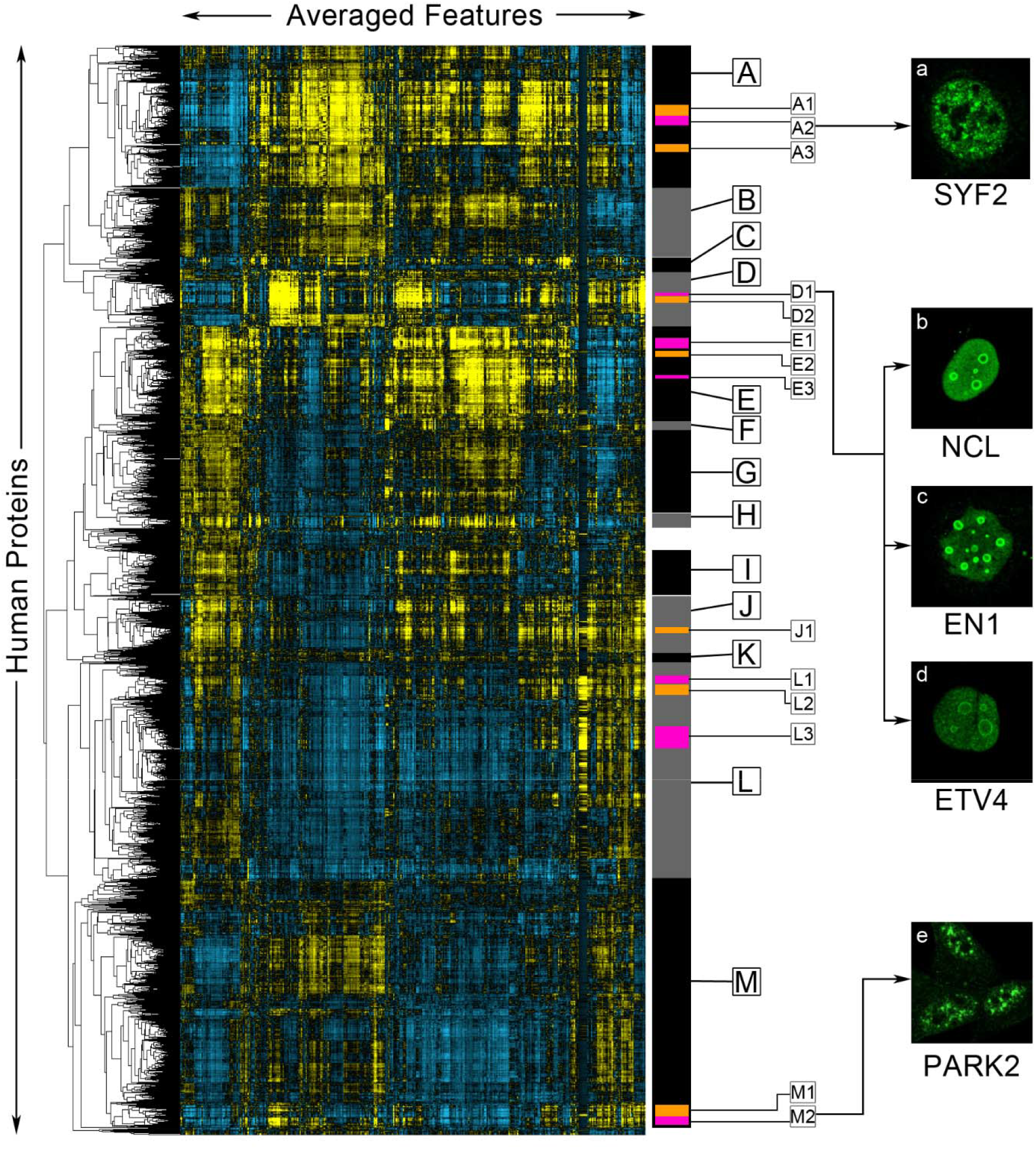
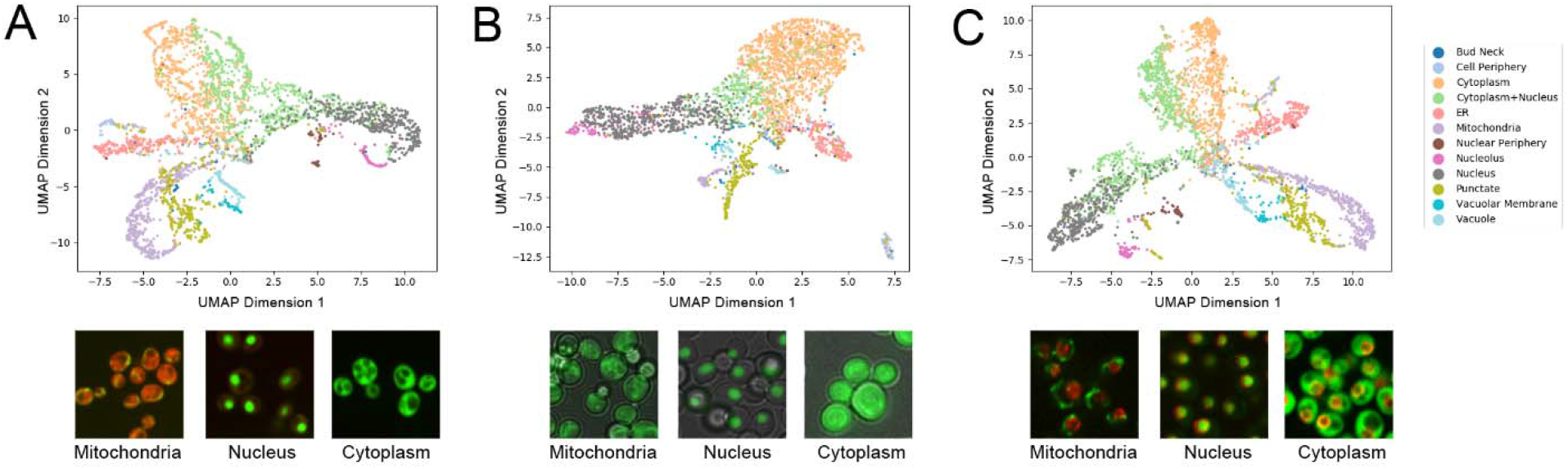
UMAP representations of protein-level paired cell inpainting representations for three independent yeast image datasets. Protein-level representations are produced by averaging the features for all single cells for each protein. A) shows the CyCLOPS WT2 dataset, B) shows the dataset of Weill *et al.*, and C) shows the dataset of Tkach *et al.* All UMAPs are generated with the same parameters (Euclidean distance, 20 neighbors, minimum distance of 0.3). Embedded points are visualized as a scatterplot and are colored according to their label, as shown in the shared legend to the right; to reduce clutter, we only show a subset of the protein localization classes. Under each UMAP, we show representative images of mitochondrial, nucleus, and cytoplasm labels from each dataset.

### Human cells with similar proteins are closer in the paired cell inpainting feature space

To further test the generalizability of the method, we trained a model for human fluorescent microscopy images with paired cell inpainting, on the entirety of the Human Protein Atlas, encompassing 638,640 single cells from 41,285 images. While the Human Protein Atlas does contain information about the general protein localization for each image, our goal is to demonstrate that our method can work on high-content human datasets without expert labels. Hence, for training our model, we do not make use of these labels in any form.

To evaluate the feature representation of single human cells, we directly analyzed the pairwise distances in the feature space. In contrast to the previous analysis (Figure 2A), we do not have a labeled single cell dataset for human cells: annotations in the Human Protein Atlas are at image-level. We therefore did not analyze classification performance. Instead, since we expect features to give similar representations to cells with similar protein localization patterns (ideally independent of cell morphologies, illumination effects, or cell lines), we compared the distance between cells with similar localization patterns in the feature space to cells with different localization patterns. We computed the normalized average distance between cells in images for proteins with the same gene ontology (GO) localization term (see Methods): a more negative score indicates that cells with the same GO term are on average closer together in the feature space (Figure 2B). We observed that features from both the Conv3 (−0.574 overall) and Conv4 (−0.349 overall) layers show smaller distances for proteins with the same GO localizations than classic computer vision features (−0.274 overall), autoencoder features (−0.281 overall), and features transferred from a pre-trained VGG16 model (−0.325 overall).

To test the robustness of the feature representation to morphological differences, we repeated the above experiment, but controlling for cell line. We reasoned that if a feature set was robust, cells with similar localization patterns would show smaller distances, even if the cells had different morphologies. We compared the normalized average pairwise distance between cells from images with the same GO localization term, when both images were constrained to be from different cell lines, compared to when both images were constrained to be from the same cell lines (Table 2). We note that we normalized the results shown in Table 2 differently from that of Figure 2B, so that the distances are not directly comparable (see Methods).

**Table 2.** Normalized average pairwise distances between cells with the same GO localization term in various feature spaces.

| Feature Set | Normalized Average Pairwise Distance (Same Cell Lines) | Normalized Average Pairwise Distance (Different Cell Lines) | Difference |
| --- | --- | --- | --- |
| **Classic** | -0.3027 | -0.1813 | 0.1214 |
| **Transfer Learning (VGG16)** | -0.2525 | -0.1515 | 0.1010 |
| **Autoencoder** | -0.2486 | -0.0965 | 0.1521 |
| **Paired Cell Inpainting (Conv3)** | **-0.4855** | **-0.3953** | **0.0902** |
| **Paired Cell Inpainting (Conv4)** | **-0.2656** | **-0.2271** | **0.0385** |

Normalized average pairwise distance scores between cells in image pairs with the same GO localization term, with the additional constraint of both images between from either the same cell line, or from different cell lines. We also report the difference between the two sets of scores; a smaller difference means that the model’s performance drops less when cell lines are different.

As expected, all feature sets have higher distances in their feature spaces for cells with similar localizations, when cells come from different cell lines. However, paired cell inpainting features have the smallest drop, and still perform the best of all feature sets, even when cells come from different cell lines with different typical cell morphologies. These results indicate that our method produces single cell features that not only place similar cells closer together in the feature space, but that they are more robust to morphological variation.

Interestingly, while features from Conv4 were worse at separating different protein localizations than Conv3, its features were the most robust to cell line. Based upon this observation, we hypothesize that the earlier layers in our model may represent sensitive information directly extracted from the images, whereas later layers assemble this information into a more robust, morphology-invariant form. We also noticed that the best overall performing layer differed from our yeast model to our human model. These results suggest that the behavior of layers may differ from dataset to dataset, and a preliminary evaluation of layer-by-layer properties is important for selecting the best layer for subsequent applications of the features. However, the middle layers in our model (Conv3 and Conv4) still outperformed other unsupervised feature sets in both models, suggesting that the selection of layer to use is still robust even in the absence of this preliminary evaluation.

### Proteome-wide clustering of human proteins with paired cell inpainting features discover localization classes not visible by human eye

We next sought to demonstrate the utility of the features to various applications in cell biology. First, we used the paired cell inpainting features from Conv3 of our human model in an exploratory cluster analysis of human proteins (Figure 4). Similar clustering experiments have been performed for yeast proteins, and have found clusters that not only recapitulate previously-defined subcellular localization patterns, but discover protein localization and function at a higher resolution than manual assignment (1). For this analysis, we included every protein in the Human Protein Atlas that we obtained enough single cells for (a total of 11995 of 12068 proteins.) All of these proteins are shown in Figure 4. We averaged the features of all single cells for each protein (pooling together all single cells from different cell lines and antibodies), and clustered the averaged features using hierarchical agglomerative clustering. While hierarchical clustering has previously been applied to a subset of human proteins in a single cell line to identify misannotated proteins (49), our analysis is, to our knowledge, the most comprehensive for the Human Protein Atlas.

**Fig 4.**
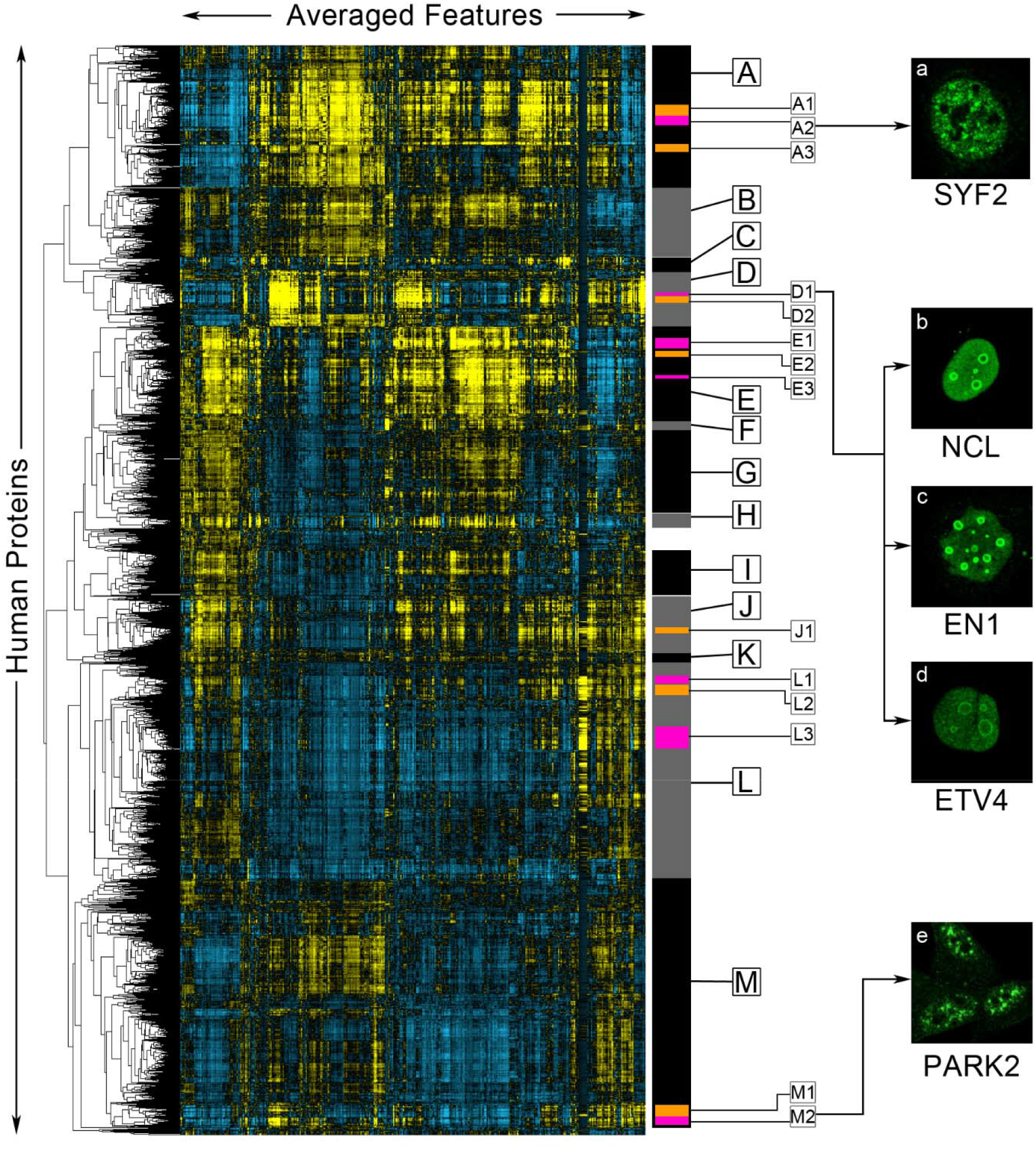
A clustered heat map of paired cell inpainting feature representation of proteins in the Human Protein Atlas. Proteins are ordered using maximum likelihood agglomerative hierarchical clustering. We visualize the average of paired cell inpainting features for all cells for each protein as a heat map, where positive values are colored yellow and negative values are colored blue, with the intensity of the color corresponding to magnitude. Columns in this heat map are features, while rows are proteins. Features have been mean-centered and normalized. We manually select major clusters (grey and black bars on the right), as well as some sub-clusters within these major clusters (orange and purple bars on the right), for enrichment analysis, presented in Table 3. On the right panels (panels a-e), we show representative examples of the protein channel (green) of cells from proteins (as labeled) in the clusters indicated by arrows from the images.

**Table 3.**
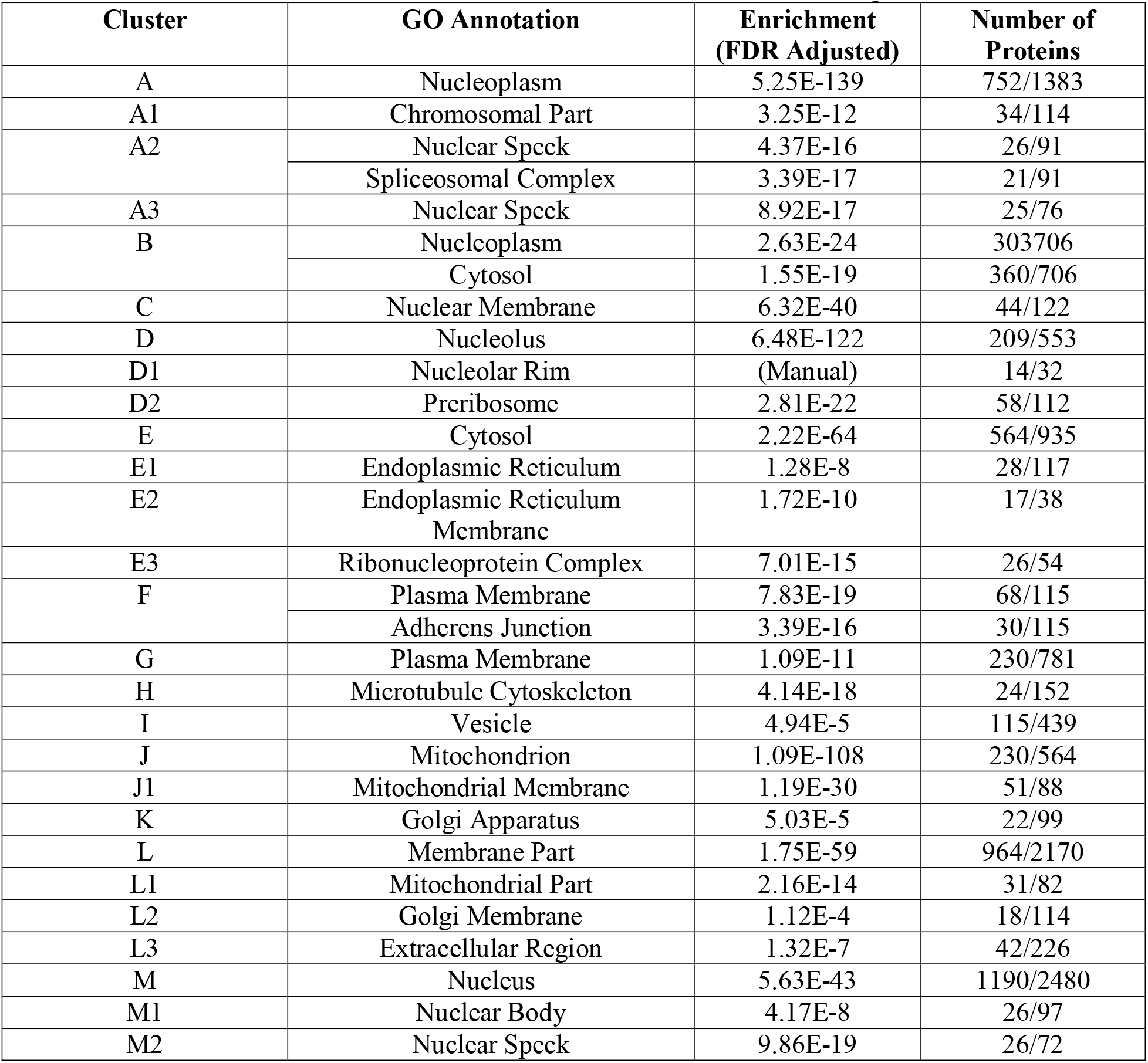
Enrichments for clusters identified in Figure 4.

Gene ontology (GO) enrichments for manually identified clusters in Figure 4. For each cluster, we report selected GO annotation terms in the component ontology for *Homo sapiens*, their enrichment (adjusted with the false discovery rate), and the number of proteins with the enrichment versus the total number of proteins in the cluster. We used all human proteins in our clustering solution as the background for these enrichments. We used GOrilla (50), a bioinformatics tool for identifying enriched GO terms. Full lists of the proteins in each cluster are available as part of Supplementary Data 1.

We find that proteins in the same cellular components generally cluster together (Table 3). We identified clusters enriched for about 15 of the 18 localization classes with more than 30 proteins. As these are the cellular components manually annotated by the Human Protein Atlas, these results suggest that our unbiased, unsupervised clustering of proteins broadly agrees with the protein localization classes assigned by biologists. Within large clusters of more general localizations, we also identified smaller sub-clusters of protein complexes or functional protein classes, including the spliceosomal complex (Cluster A2), the ribonucleoprotein complex (Cluster E3), and the preribosome (Cluster D2). These are cellular components that are not typically identifiable by human eye, suggesting that our features are describing the localization of the proteins at a higher-resolution than human annotation.

To assess the ability of our clustering to find rare localization patterns, we asked if we could identify proteins localized to the nucleolar rim. Out of 226,732 images in the Human Protein Atlas, Sullivan *et al.* only classify 401, or 0.18%, as nucleolar rim (20); based on this proportion, we estimated that ~20 proteins in our clustering solution would localize in this manner. The version of the Human Protein Atlas used in this study did not incorporate the nucleolar rim annotation, but the documentation gives one example, MKI67 (24). We found MKI67 in Cluster D1. Of the 32 proteins closest to MKI67 in our hierarchical clustering, we identified 13 other proteins with an unambiguous nucleolar rim localization (we list these proteins in Supplementary Data 1 and provide links to their images in the Human Protein Atlas), although there may also be others too ambiguous to determine by eye. We show three of these in Figure 4, NCL (Figure 4b), EN1 (Figure 4c), and ETV4 (Figure 4d). This supports the idea that unsupervised analysis in the feature space can identify rare patterns for which training classifiers would be difficult.

We observed several clusters of nuclear speck proteins, including clusters A1, A2, and M2. We decided to evaluate images for proteins in these clusters to determine if there is any distinction in their phenotype. We observed that proteins in clusters A1 and A2 tended to localize purely to nuclear specks (Figure 4a), whereas proteins in cluster M2 tended to localize to both nuclear specks and the cytosol (Figure 4e). Consistent with this interpretation, we found a strong enrichment for the spliceosomal complex for cluster A1, but no enrichment for cluster M2. To further test the possibility that the unsupervised analysis in the feature space reveals higher resolution functional information than expert annotation, we asked if our features could be used to discover higher-resolution subclasses of proteins that are difficult for experts to annotate visually. First, previous work with vesicle-localized proteins in the Human Protein Atlas has suggested that these proteins can be distinguished into many subclasses (2,51). We clustered features for human proteins annotated to localize to the vesicle only, and found structure in the feature representation that corresponds to visually-distinct vesicle patterns (Abundantly and more evenly distributed in the cytosol, concentrated closer to the nucleus or sparse puncta, Supplementary Figure 2, Clusters A, B and C respectively). However, we were unable to associate these with any biological functions, so it is unclear if these distinctions in the patterns of vesicle distribution are functional. Next, we used features obtained from paired cell inpainting to organize proteins labelled as “punctate” in yeast dataset of images of a NOP1pr-GFP library (47) (see Methods). We find that we can distinguish Golgi, peroxisome and another cluster that contained singular or sparse foci in the cells (Supplementary Figure 4, Cluster B, C and A respectively). These analyses support the idea that unsupervised analysis of our features can reveal higher biological resolution than visual inspection of images.

### Paired cell inpainting features enable quantitative analysis of multi-localizing and spatially variable proteins

Next, we asked if features from our models could be used for the analysis of single cells of multiply-localized proteins. While multiply-localized proteins are critical to study as they are often hubs for protein-protein interactions (52,53), these proteins have posed a challenge for imaging methods. Most supervised methods remove multiply-localized proteins from their training and evaluation sets (6,9,41,54), focusing on only proteins localized to a single subcellular compartment. In supervised efforts, Sullivan *et al.* propose training with multi-label data (20), but curating this training data is challenging due to sparsity in some combinatorial classes and difficulty in manually annotating complex mixtures of patterns. In unsupervised work, unmixing algorithms (55) and generative models (56,57) have been proposed, but these methods generally rely on elaborate user-defined models of the conditional dependencies between different compartments. Thus, extending models to multiply-localizing proteins is non-trivial and an ongoing area of research.

We applied our features to analyze human single cells in images annotated to localize to the cytosol-and− nucleoplasm and to the nucleus-and-nucleoli by the Human Protein Atlas. Using the single-cell features from Conv3 of our human model, we measured a simple score based on distances in our feature space (see Methods). For our cytosol-and-nucleoplasm analysis, a negative score indicates that the cell is closer to the average cytosol-only cell than the average nucleoplasm-only cell in the feature space, while a positive score indicates that the cell is closer to the average nucleoplasm-only cell than the average cytosol-only cell. For our nucleoli-and-nucleus analysis, a negative score indicates that the cell is closer to the average nucleoli cell than the average nucleus cell, while a positive score indicates that the cell is closer to the average nucleus cell than the average nucleoli cell. We calculated this score for every single-cell localized to both compartments, or to either compartment alone (Figure 5).

**Fig 5.**
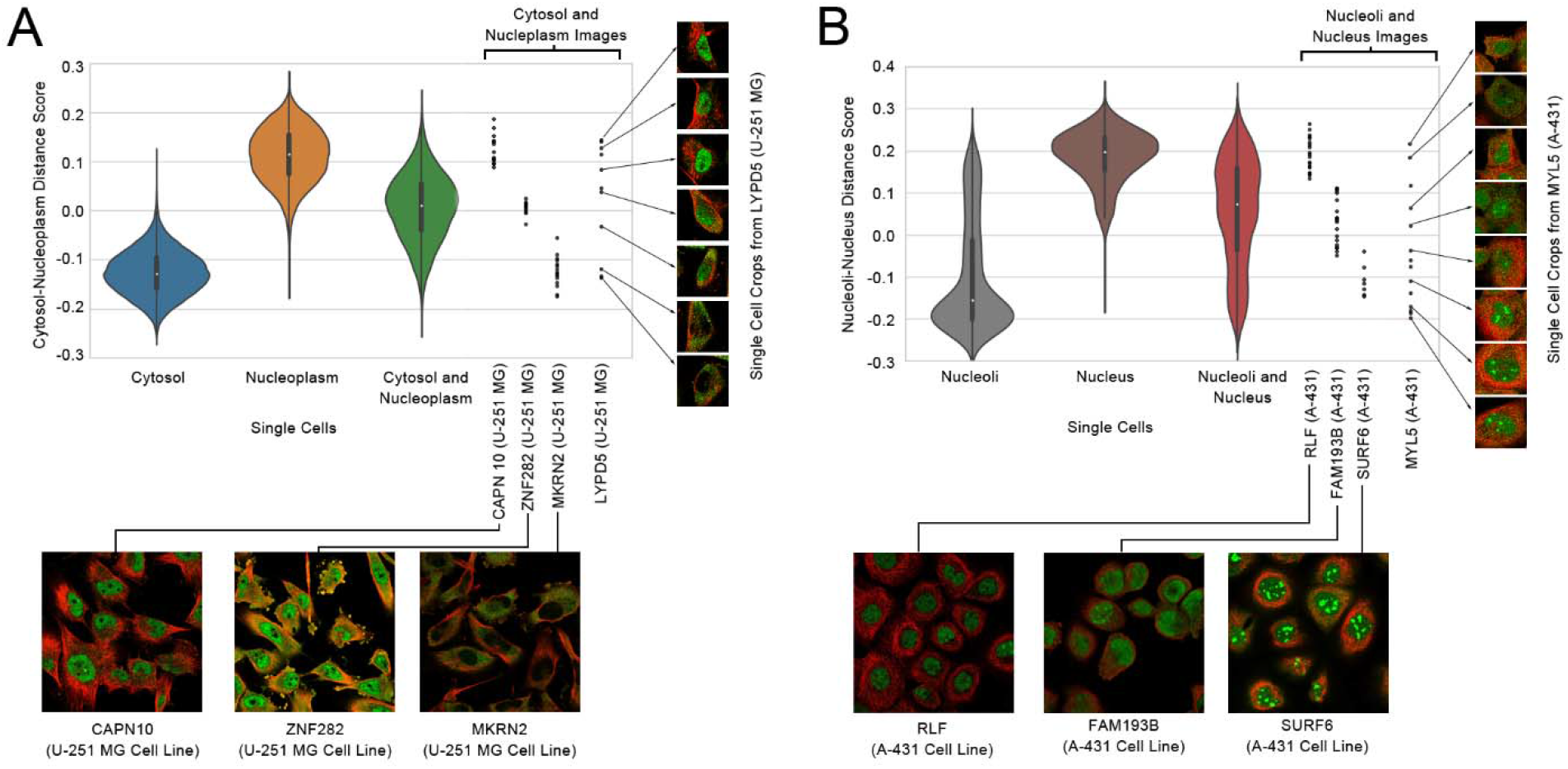
Single-cell scores (as explained in the text) for cytosolic-and-nucleoplasm cells (A), and nucleoli-and-nucleus cells (B). We visualize single cells as violin plots. The black boxes inside the violin plots show the interquartile ranges and the white dots show the medians. To the right of the violin plots, we show all of the single-cells for four different images of proteins annotated to be multi-localizing to either the cytosol-or-nucleoplasm or the nucleolus-and-nucleus as black dots, as labeled in the x-axis. For each image, we show either a representative crop of the image, or multiple single cell crops; in both cases, the protein is shown in green, while the microtubule is shown as red. Note that while the violin plots include all cell-lines, we show four examples from the same cell-line for each plot, to demonstrate that images from the same cell-line can have different scores depending on their phenotype.

We observe that single-cells with a cytosol-only annotation have a more positive score, while single-cells with a nucleus-only annotation have a more negative score (Figure 5A). In contrast, single-cells annotated to localize to both compartments have a score closer to 0. We observed that the skew of multiply-localized single-cells towards a positive or negative score reflects the proportion of protein localized to the two compartments: for example, for single cells stained for the spatially-variable protein LYPD5 in the U-251 MG cell line, we observed that positive scores corresponded to a more dominant nuclear localization, negative scores corresponded to a more dominant cytosol localization, and scores closer to 0 corresponded to a more even mix of both localizations (Figure 5A). We made similar observations for nucleoli-and-nucleus single-cells (Figure 5B). However, we also noticed that while most of the single-cells with a nucleoli-only annotation had negative scores, some had more positive scores, suggesting that the nucleoli-and-nucleus images are often annotated as nucleoli-only. These results suggest that simple distances in our feature space could be used to not only identify multiply-localized proteins, but also quantitate the relative distribution of protein between its compartments. We have included the mean scores for every cytosol-and-nucleoplasm and nucleoli-and-nucleus protein by cell line as Supplementary Data 2.

We observed that proteins with spatial-variability sometimes had different distributions over our scores compared to more homogeneously-localized proteins (Figure 5). This observation suggested that the standard deviation could be useful in discovering proteins with single-cell variability. We looked at the 10 images with the highest standard deviations for cytosol-and-nucleoplasm single-cells, and for nucleoli- and-nucleus single-cells. For cytosol-and-nucleoplasm single-cells, 6/10 are previously annotated by the Human Protein Atlas to have single-cell variability, and we observed unambiguous, but previously unannotated, single-cell variability in the images for the remaining 4. For nucleus-and-nucleoli single-cells, 5/10 are previously annotated, and we observed unambiguous and previously unannotated single-cell variability for the remaining 5 (we show one example of a previously unannotated image in Figure 4B, for MYL5 in the A-431 cell line). While this simple metric would be less sensitive to variable proteins where the single-cells are imbalanced in their distribution of the different phenotypes, these results suggest that simple statistics in our feature space can identify at least some kinds of single-cell variability. We include the standard deviation of scores for every cytosol-and-nucleoplasm and nucleoli- and-nucleus protein by cell line in Supplementary Data 2.

### Clustering of single cells organizes spatial variability in proteins

To organize single-cell variability, without prior knowledge of the specific compartments the protein is localized to, we decided to analyze single-cells using clustering in the feature space. We selected three proteins with spatial variability at the single-cell level: SMPDL3A, which localizes to the nucleoli and mitochondria (Figure 6A); DECR1, which localizes to the cytosol and mitochondria (Figure 6B); and NEK1, which localizes to the nucleoli and nuclear membrane (Figure 6C). We clustered the single cells of each protein using hierarchical agglomerative clustering. Figure 6 shows the dendograms of the clustering solution for each protein, as well as the single-cell image crops associated with representative branches of the dendogram.

**Fig 6.**
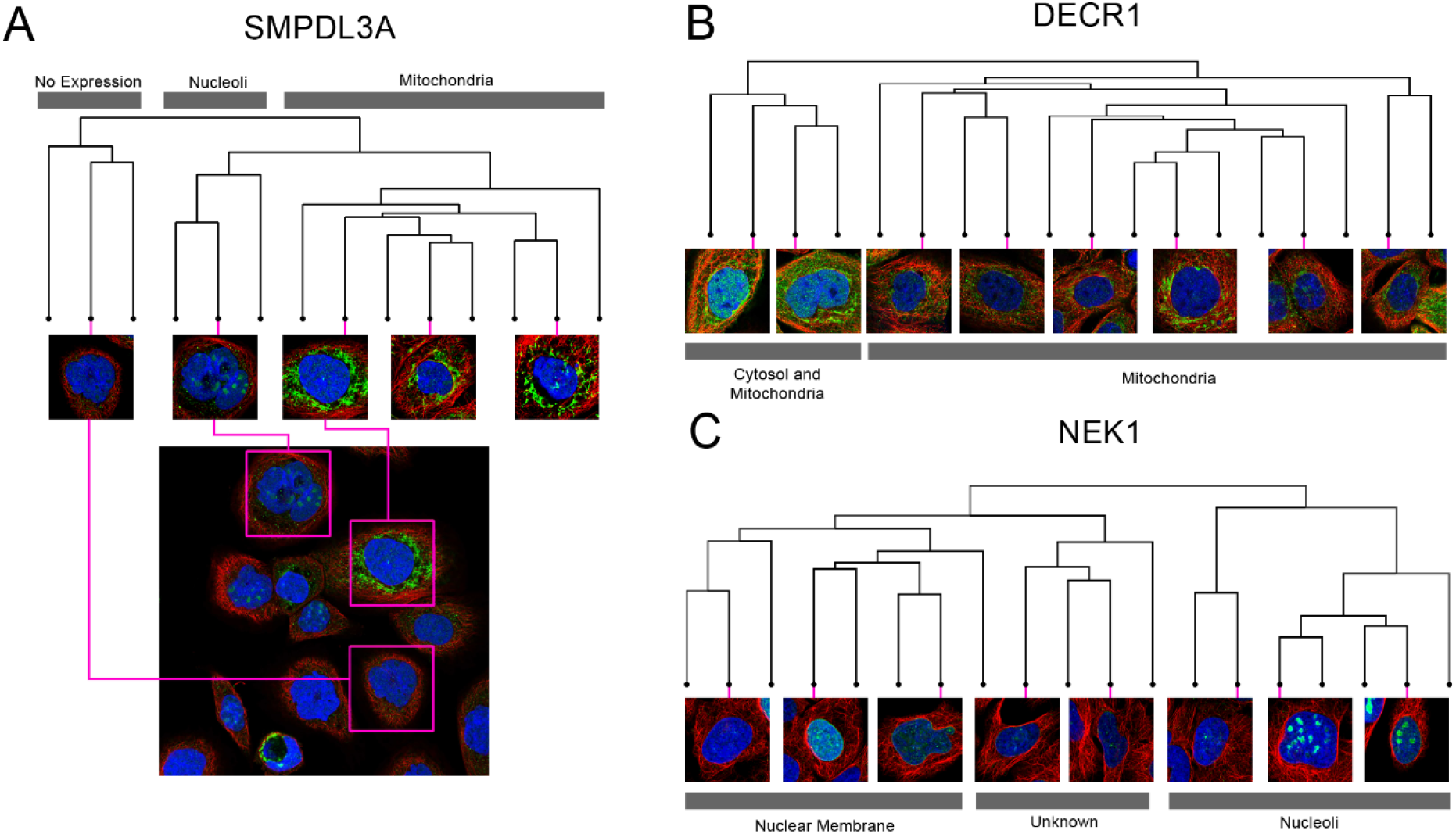
Dendograms from clustering the single-cell paired cell inpainting features of three spatially variable proteins: (A) SMPDL3A, (B) DECR1, (C) NEK1. For each dendogram, we show single-cell image crops associated with representative branches of the dendogram. For (A) SMPDL3A, we also show a field of view of the image, and where various single-cell crops originate from in the full image (pink boxes). We label clades in the dendogram (grey boxes) with the apparent localization of the single cells.

In general, we observe distinct clusters of single cells for spatially variable proteins, visible as distinct clades in the dendograms (Figure 6). For SMPDL3A, we observe clusters of mitochondria-localized and of nucleoli-localized single cells, and for DECR1, we observe clusters of mitochondria-localized and cytosol-and-mitochondria-localized single cells, consistent with their labels in the Human Protein Atlas. For NEK1, we observed distinct clusters for the nuclear membrane and nucleoli, consistent with its label, but also for single-cells of a third phenotype, which appeared as fine and diffuse clumps of protein within the nucleus, not recorded in the Human Protein Atlas (24). We hypothesize this may be due to the challenges in manually annotating a visually complex image with a heterogenous mixture of cells. These results suggest that single-cell clustering in the feature space can organize the phenotypes in a heterogeneous image and quantify the relative proportion of each phenotype.

## Discussion

We present a simple, but effective self-supervised learning task for learning feature representations of single cells in fluorescent microscopy images. Features learned through paired cell inpainting easily achieve state-of-the-art performance in unsupervised benchmarks for both yeast and human cells. We believe that this effectiveness is because our pretext training task controls for nuisance variation. Because our task requires the network to predict the phenotype in a different cell, the network must ignore cell-specific variation to learn more general features about the phenotype. As we show the network the target cell structural markers during training, the network is directly given experimental effects like local illumination, meaning that it does not have to learn features for these in the source cell encoder.

A key advantage to self-supervised methods is that they learn features directly from data. Our method exploits the natural structure of many imaging experiments and can be applied to learn single-cell representations for virtually any multi-channel microscopy image dataset. To demonstrate the generality of paired cell inpainting, we learned representations for four independent image datasets in this study. Two of the yeast datasets have previously only been analyzed through laborious manual inspection of thousands of images (38,47). We believe this is likely due to a lack of generality in previous automated methods: both of the classic computer vision methods we applied on the CyCLOPS dataset rely on accurate single-cell segmentation (1,58) which is challenging on the NOP1-ORF library due to highly overlapping and clumped cells, and impossible for the Nup49-RFP GFP-ORF dataset due to a lack of any cytoplasmic indicator. Similarly, neither dataset has a labeled single-cell dataset associated with it, preventing the application of supervised methods until this laborious work is complete. These datasets demonstrate important points about the context our proposed method operates in: while we show we can achieve performance gains on well-studied datasets like the CyCLOPS dataset, many biological imaging datasets are not as amenable for automated analysis. We believe that for these kinds of datasets, our method can produce similarly discriminative representations, at least on the protein level (Figure 3), enabling the fully automated analysis of these datasets without accurate segmentation or expert image labeling.

For this reason, we believe that self-supervised learning is an effective way to handle the increasing volume and diversity of image datasets as high-content imaging continues to accelerate, especially with the development of scalable tagging technologies and new libraries (47,59). To address this demand, other self-supervised methods are also emerging: Caicedo *et al.* propose training a CNN to classify cells by their drug treatment (19). We note that in contrast to their proposed proxy task, our task is predictive, rather than discriminative, and has some technical advantages in theory: since we do not force the CNN to discriminate between experiments that yield identical phenotypes, we expect that our method is less susceptible to overfitting on experimental variation. In practice, both methods work well on for their respective benchmarked applications. The choice of proxy task may be therefore be constrained by practical considerations: for example, our task cannot be applied on single-channel images, or images where cell centers cannot be automatically identified. While self-supervised learning methods for biological images are still emerging, in the future, a comparative analysis of the performance of different methods on different applications would be highly informative to researchers deciding which proxy task to apply to their own data.

While the quality of the features we learned were generally robust to different parameters and preprocessing operations on the dataset, outperforming other unsupervised methods regardless of settings, we found several factors that we believe could improve the features. First, filtering proteins that have single-cell variability in their protein expression, for example, by using labels in the Human Protein Atlas (24). Likely, this would reduce amount of noise caused by pairing single cells with different protein expression during training. Second, while our method does not rely on accurate single-cell segmentations (1,40), it uses a crop centered around single cells in the image. We found that the method we used to detect cell centers for cropping was important, with classic segmentation techniques based on watershedding (1) resulting in poorer features the methods used here (see Methods). We hypothesize that this is because artifacts included as cells increase noise during training when a cell gets paired with an artifact.

Here, we focused on the proxy task. Like most previous self-supervised work, we used a simple AlexNet architecture (17–19,29,30). However, future optimizations to the architecture will likely improve the applicability and performance of our method. A major limitation of self-supervised methods that use the AlexNet architecture is that the layer to extract features with must be determined. For most proxy tasks, the best performing-layer is an intermediate convolutional layer in AlexNet (17–19,29,30). However, recent work optimizing architectures for self-supervised methods suggests that architectures with skip-connections can prevent the quality of self-supervised representations from degrading in later layers (60). Applying these insights to our method might mean that the final layer could always be chosen for transfer to new tasks, improving the practicality of the method.

We note that some technical points specific to our method may also improve performance. First, some proteins localize in a pattern that cannot be predicted deterministically from the structural markers of the cell alone. The inability of the network to accurately reconstruct these proteins may limit its feature learning. Future work in loss functions that encourage more realistic outputs, such as adversarial losses, could improve this issue. Second, for our experiments, we iterated over all cells in our training dataset and sampled pairs randomly. A more sophisticated pair sampling scheme may improve the quality of the features: for example, focusing on harder pairs where the morphology of the cells differs more drastically. There are also some protein localization patterns correlated with the cell cycle, such as the mitotic spindle. Due to their low penetrance in an image (for example, only a fraction of the cells will be undergoing mitosis), we do not expect our model to learn these patterns very well. Even if these cells are paired correctly, there may only be one or two cells with the specific cell cycle stage required to exhibit the pattern in the entire image. Better datasets with more fields of view, or work in finding an unsupervised method for balancing the patterns in the data, may improve this issue.

In the final aspect of this work, we performed, to our knowledge, the most comprehensive unsupervised analysis of protein localization spanning the entire Human Protein Atlas to date. Despite pooling all cell lines and antibodies together for proteins, we capture enrichments for high-resolution cellular components in our cluster analysis. Moreover, we showed that we could identify rare phenotypes, and use our features to discover distinct subclasses of proteins unsupervised. These results emphasize the importance of unbiased analysis, in contrast to supervised approaches. Rare phenotypes may be excluded from classifiers due to lack of training data, or lack of knowledge of which images to mine training data from. We discovered several proteins with a nucleolar rim localization, based upon the one example given by the Human Protein Atlas. Similarly, pre-defined classes may hide functionally-important variability within classes, and may be biased by human pre-conceptions of the data. We showed that coarse human-annotated classes can be clustered into distinct subclasses, and that functional protein classes such as the preribosome or splicesomal complex are clustered in protein localization data. Overall, these results demonstrate the power of unsupervised analysis in discovering new and unexpected biology, a task that supervised methods are fundamentally unable to perform, no matter how accurate they are at annotating pre-defined human knowledge.

## Code Availability

Code and pre-trained weights for the models used in this work are available at https://github.com/alexxijielu/paired_cell_inpainting.

## Acknowledgements

We thank Shadi Zabad, Taraneh Zarin, Nirvana Nursimulu, and Purnima Kompella for valuable comments on the manuscript. We thank Brenda Andrews and Helena Friesen for help in accessing the CyCLOPS data. We thank Maya Schuldiner, Uri Weill, and Amir Fadel for help in accessing the NOP1pr-ORF yeast collection. We thank Grant Brown and Brandon Ho for help in accessing the Nup49-RFP GFP-ORF yeast collection. This work was conducted on a GPU generously provided by Nvidia through their academic seeding grant. This work was funded by the National Science and Engineering Research Council (Pre-Doctoral Award), Canada Research Chairs (Tier II Chair), and the Canadian Foundation for Innovation.

---

**Fig S1.**
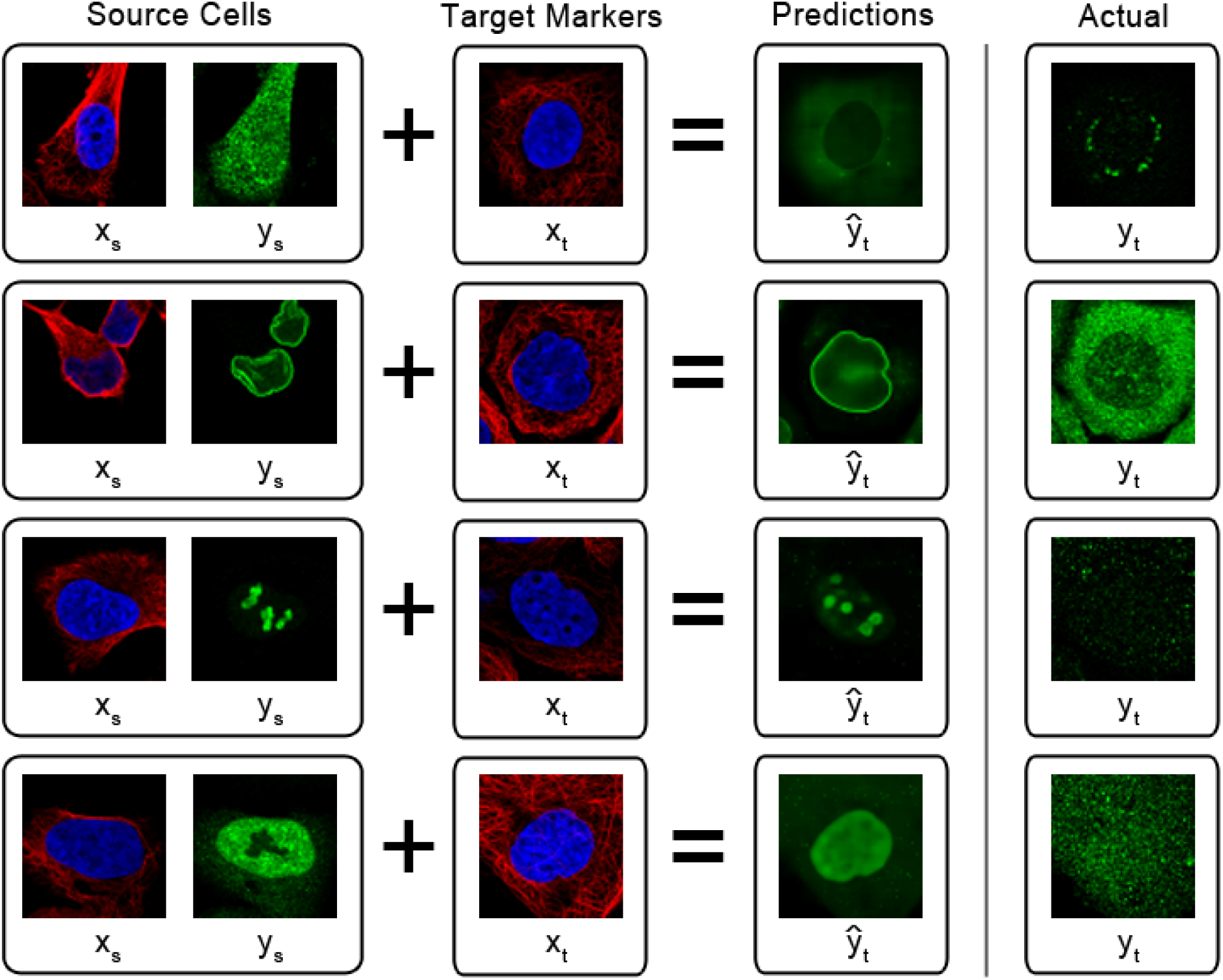
End-to-end paired cell inpainting results for pairs of cells unseen during training. We take four source human cells with different localizations (left-most boxes), belonging in the cytosol, nuclear membrane, nucleoli, and nucleoplasm, from top to bottom. For each of our source cells, we run our human cell model end-to-end with target markers from different images (left-center boxes) as input to generate predictions of the protein localization in these target cells (right-center boxes). We show the actual protein localization of the target cells (right-most boxes) to demonstrate that our model is capable of synthesizing results based on the source cells even when there is a mismatch between the source and target cell protein localization.

**Fig S2.**
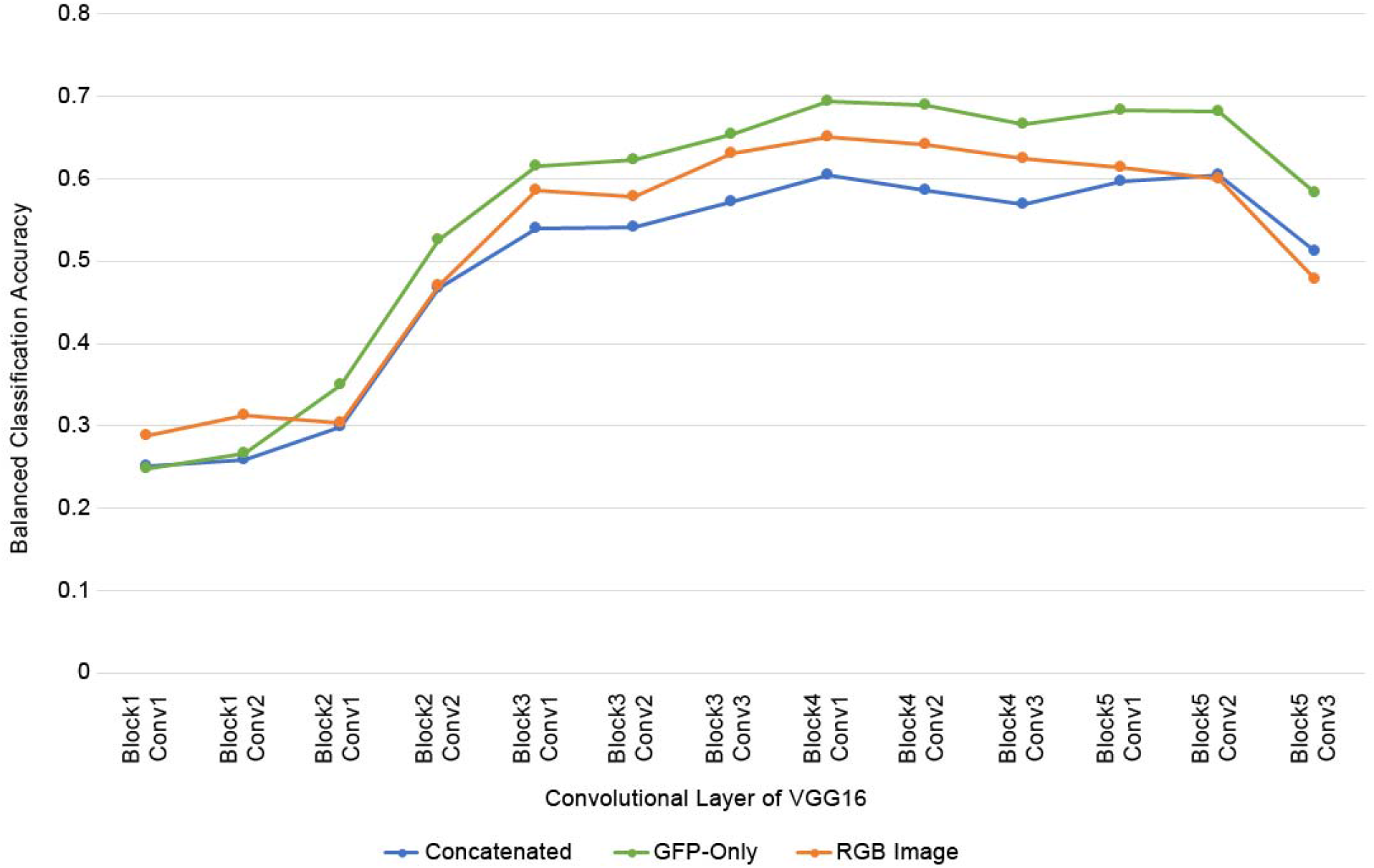
Classification accuracies for representations extracted over each convolutional layer of a VGG16 model pretrained on ImageNet, for different input strategies. Images in the yeast single cell classification dataset were inputted using three different strategies: for “Concatenated”, we inputted the RFP and GFP channels independently as greyscale images and concatenated the representations; for “GFP-Only”, we inputted the GFP channel as a greyscale image and used this representation alone; and for “RGB Image”, we arbitrarily mapped channels to RGB channels (RFP to red, GFP to green, and blue left empty). Feature representations were extracted by maximum pooling the feature maps over spatial dimensions. We report the balanced classification accuracy using a leave-one-out kNN classifier (*k* = 11) for these representations, identical to the one described in the “Paired cell inpainting features discriminate protein subcellular localization in yeast single cells” section of the Results.

**Table S1.**
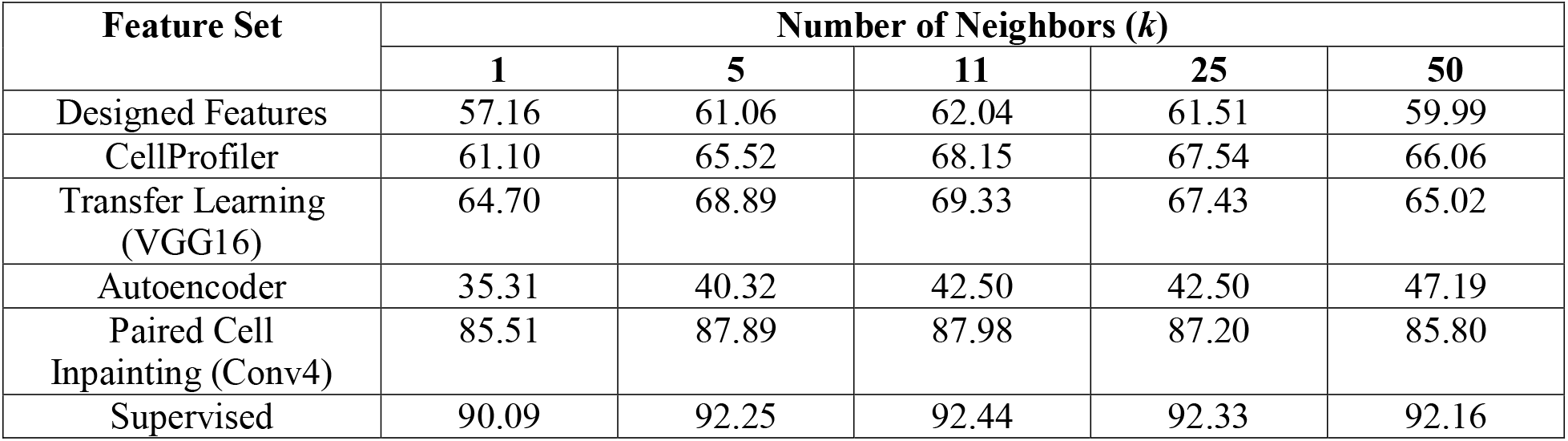
Classification accuracies for feature sets with various parameterizations of *k*.

| Feature Set | Number of Neighbors ($k$) | | | | |
| --- | --- | --- | --- | --- | --- |
|  | 1 | 5 | 11 | 25 | 50 |
| Designed Features | 57.16 | 61.06 | 62.04 | 61.51 | 59.99 |
| CellProfiler | 61.10 | 65.52 | 68.15 | 67.54 | 66.06 |
| Transfer Learning (VGG16) | 64.70 | 68.89 | 69.33 | 67.43 | 65.02 |
| Autoencoder | 35.31 | 40.32 | 42.50 | 42.50 | 47.19 |
| Paired Cell Inpainting (Conv4) | 85.51 | 87.89 | 87.98 | 87.20 | 85.80 |
| Supervised | 90.09 | 92.25 | 92.44 | 92.33 | 92.16 |

Classification accuracies for single yeast cell localization classes using a *k*NN classifier on our test set of 30,889 labeled single cells, using various feature representations. We report the overall accuracy as the balanced accuracy of all classes.

**Table S2.** Classification accuracies for feature sets with logistic regression and random forest classifiers.

| Feature Set | Accuracy<br>(Logistic Regression) | Accuracy<br>(Random Forest) |
| --- | --- | --- |
| Designed Features | 69.17 | 69.80 |
| CellProfiler | 80.00 | 78.17 |
| Transfer Learning<br>(VGG16) | 83.77 | 72.29 |
| Autoencoder | 63.33 | 56.41 |
| Paired Cell<br>Inpainting (Conv4) | 91.34 | 87.49 |
| Supervised | 94.05 | 92.94 |
| Supervised<br>(End-to-End) | 94.02 |  |

Classification accuracies for single yeast cell localization classes using logistic regression and random forest classifiers on our dataset of 30,889 labeled single cells, built on various feature representations. We also report accuracy of the classifier end-to-end for the fully-supervised CNN in the last row of the table. Metrics are reported as the average balanced accuracy on the test sets under 5-fold cross-validation. We implemented all classifiers in Python using the scikit-learn package. For our logistic regression classifiers, we used a L1 penalty with a balanced class weight. For our random forest classifiers, we used 500 trees with 20% of the features used to determine best splits, and a balanced class weight.

**Fig S3.**
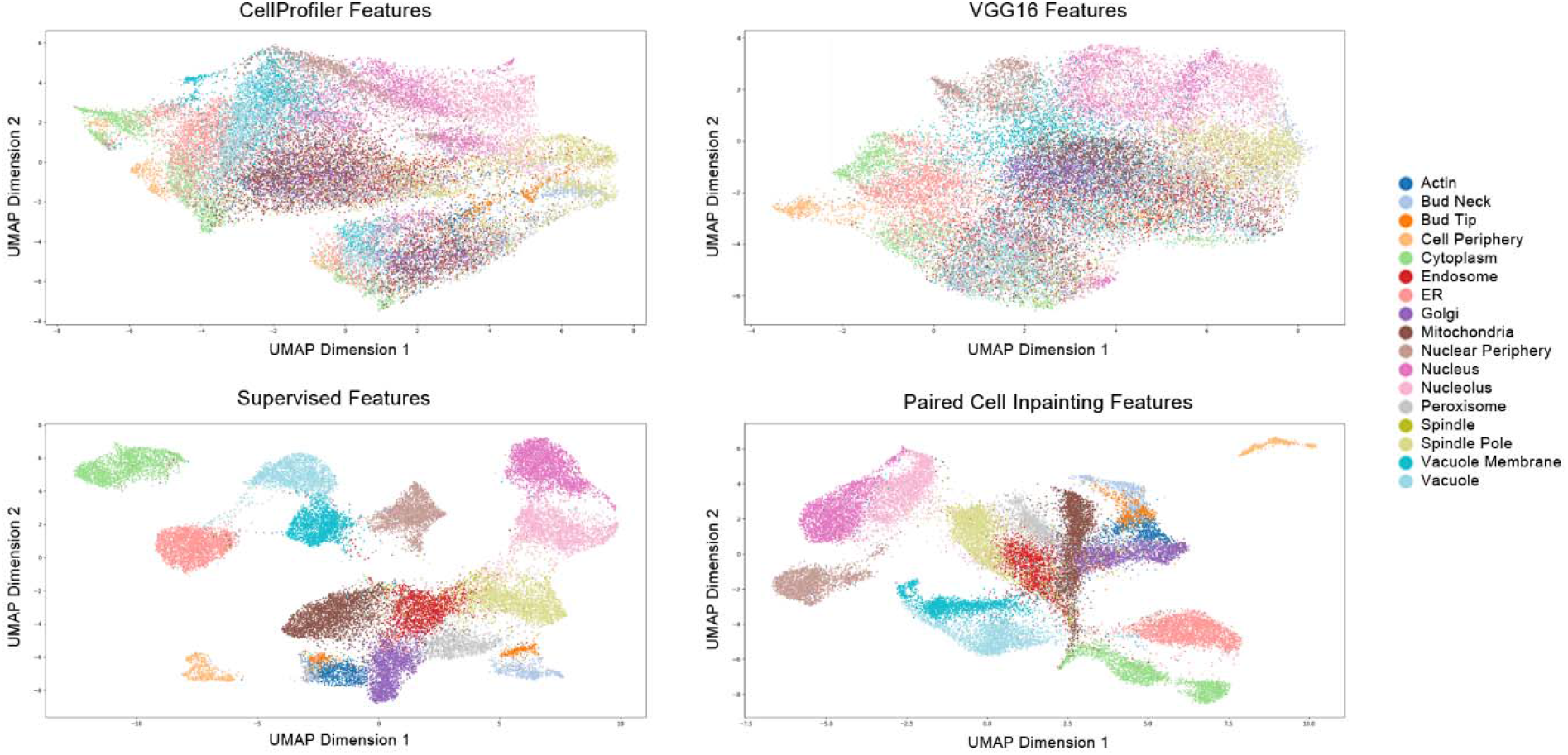
UMAP representations of various features, as labeled above each scatterplot, for our labeled single yeast cell benchmark dataset. All UMAPs are generated with the same parameters (Euclidean distance, 30 neighbors, minimum distance of 0.3). Embedded points are visualized as a scatterplot and are colored according to their label, as shown in the legend to the right.

**Fig S4.**
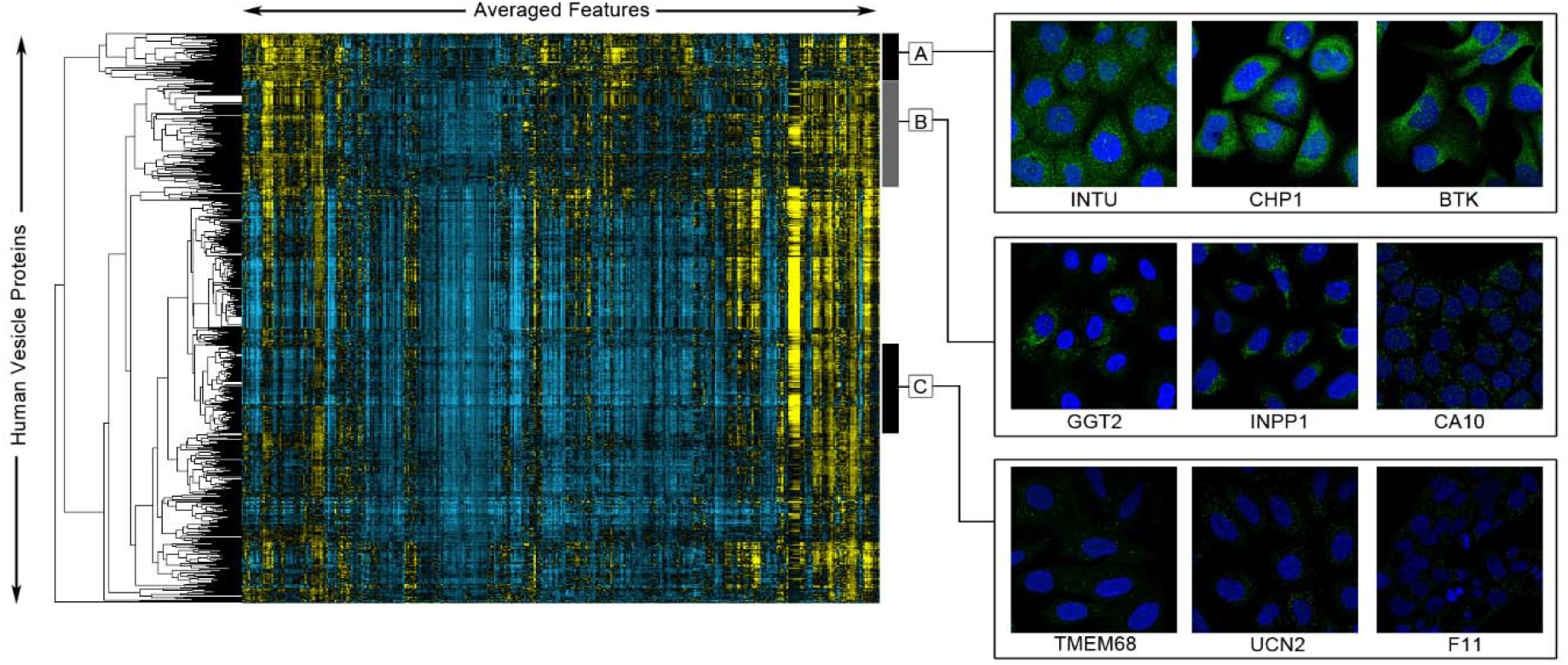
Averaged paired cell inpainting features for vesicle-only proteins in the Human Protein Atlas, using features from Conv3 of our human model trained on the Human Protein Atlas dataset, ordered using maximum likelihood agglomerative hierarchical clustering. We visualize features as a heat map, where positive values are colored yellow and negative values are colored blue, with the intensity of the color corresponding to magnitude. Columns in this heat map are features, while rows are proteins. Features have been mean-centered and normalized using all proteins in the dataset. We show three clusters (black and grey bars on the right), and crops of three representative images of the proteins within each of the clusters. For image crops, we show the protein channel in green, and the nucleus channel in blue.

**Fig S5.**
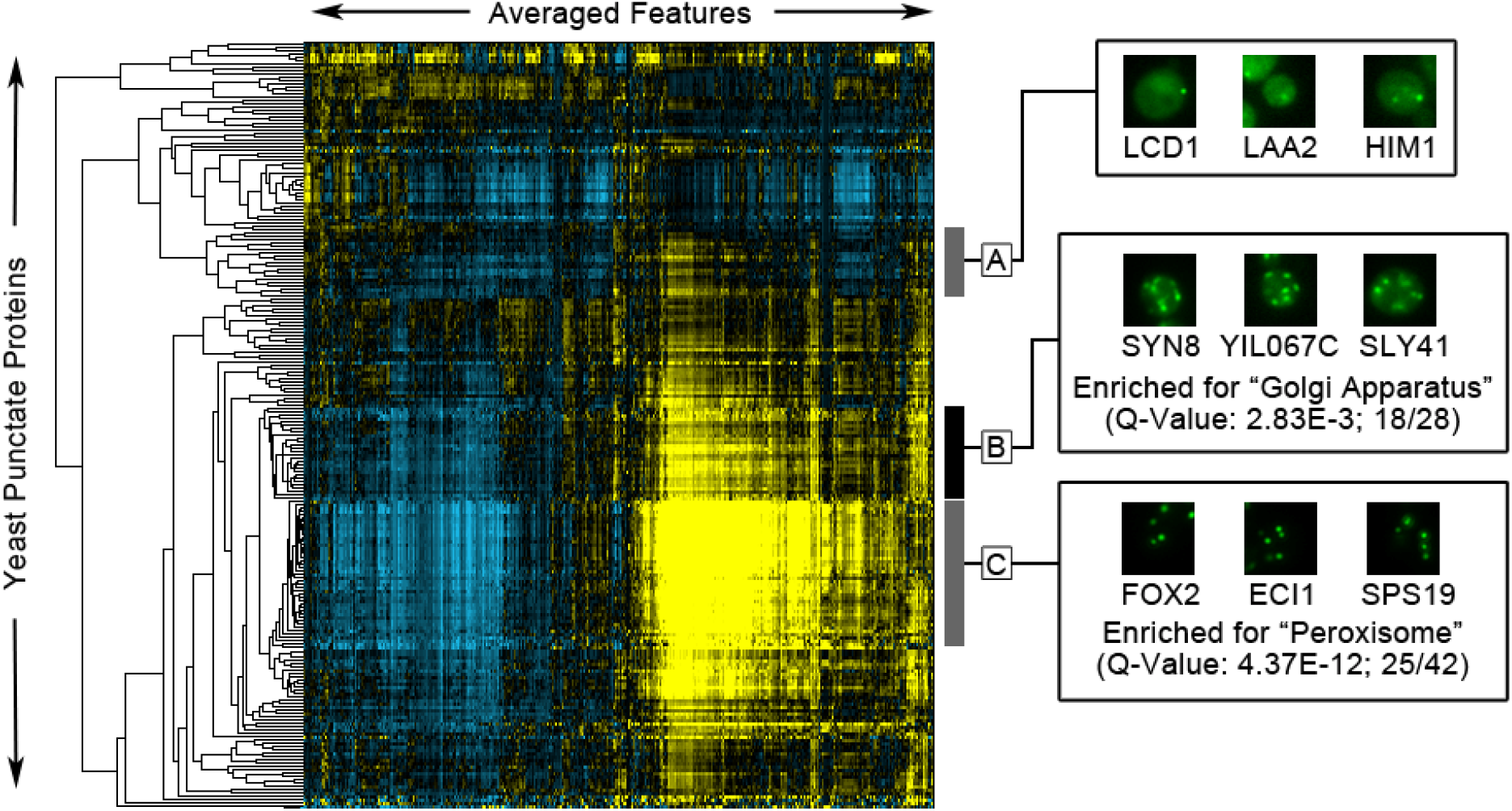
Averaged paired cell inpainting features from Conv4 of our yeast model trained on the NOP1pr-GFP dataset, for proteins labeled as punctuate in the NOP1pr-GFP library images, ordered using maximum likelihood agglomerative hierarchical clustering. We visualize features as a heat map, where positive values are colored yellow and negative values are colored blue, with the intensity of the color corresponding to magnitude. Columns in this heat map are features, while rows are proteins. Features have been mean-centered and normalized using all proteins in the dataset. We show three clusters (black and grey bars on the right), and crops of three representative images of the proteins within each of the clusters. For image crops, we show the protein channel only in green. For clusters B and C, we show the GO enrichment of the clusters relative to all punctate proteins; we list the q-value (the FDR-corrected p-value) and the number of proteins in the cluster with that annotation relative to the full size of the cluster.

